# Label-free flow operators and stability operators from single-cell time courses: what local dynamical operators can and cannot identify

**DOI:** 10.64898/2026.11.05.764643

**Authors:** David Redd, Sam Green, Tommy W. Terooatea

## Abstract

Local Jacobians of single-cell dynamics are used to locate unstable cells along trajectories, but what they identify from measured data has rarely been tested against independent truth. We present scJDO2, which separates two per-cell operators by the time information they require: a score operator, the Hessian of a score-matched log density, available from any snapshot; and a flow operator, the Jacobian of a drift fitted to a measured velocity, here estimated label-free from a sampled time course by entropic optimal transport. Both are summarized in a shared operator space of magnitude-free descriptors, with the invariance of each stated, and evaluated with pre-registered tests.

On synthetic data calibrated to six real datasets, the flow operator from the time course alone separated fate-bifurcation cells from committed cells on held-out seeds (AUC 0.89–1.00) with a-priori settings; tuning on development seeds overfit. On LARRY lineage-traced hematopoiesis, its fitted drift predicted the clonal displacement of barcoded cells beyond population movement (Spearman 0.56, permutation p < 0.001; 0.50 after removing the mean shift) and a held-out sampling day better than mean-displacement and no-movement baselines (energy distance 0.69, 95 % CI 0.66–0.73, vs 0.87 and 0.88); lineage thus validates the velocity, not λ□. Mini-batch transport reproduced dense transport (velocity cosine 0.9998) and scaled to all 130,887 cells (transport 5 min and 2.5 GB; drift fit and Jacobians 21 min and 1.3 GB). On an independent simulator it detected two of three bifurcations and two of three cycles and never reported instability on linear trajectories.

Several readings failed. Neither operator recovers the values of the true Jacobian; the flow operator’s leading eigenvalue ranks branching cells empirically. The flow operator depends on the transport metric and inverts on Factor Analysis. Operator archetypes isolated bifurcation cells but did not outperform expression clustering and were unstable across fit seeds. The score operator did not predict CRISPRi knockdown responses or sister-cell fate coherence. A confirmatory test of transition timing could not be constructed, and diffusion is not identifiable at typical sampling intervals. The released software reports these diagnostics.

**Changes from v56:**

1. **Two operators, matched to the time information measured (new).** v56 used one drift field learned by diffusion score matching (DSM) on pseudotime. v57 (scJDO2) distinguishes the **score operator** *J_score = β□² log p* (from a snapshot; it describes the curvature of the cell-state density) from the **flow operator** *J_flow = ∂f/∂z* of a drift *f* fitted to a **measured** velocity (time-course optimal transport, lineage, metabolic labeling or RNA velocity).
2. **Operator space (new); archetypes (carried over).** Both operators are summarized by magnitude-free descriptors in one operator space; spectral descriptors of the flow operator are invariant under any linear change of coordinates, two eigenvector-based descriptors only under orthogonal ones. Operator archetypes were introduced in scJDO v1 [1] and are carried over; v57 fits them in the new operator space and tests them against true regimes.
3. **Label-free OT flow operator, evaluated (new).** Flow operators are estimated from a time course alone (entropic OT between sampling days → barycentric velocity → drift → Jacobian) and evaluated on held-out seeds, an independent simulator and lineage barcodes, and scaled to a full dataset. OT velocities and Jacobians of OT-learned drifts have precedent [2–4]; the evaluation is what is new.
4. **Identifiability analysis (new):** representation, clock, rotation, stationarity, diffusion and composition, with an explicit list of what was tested and not supported.
5. **Correction:** v56 recommended Factor Analysis (FA) as the default representation. This **does not hold for flow operators**: the OT flow operator inverts on FA (fork AUC 0.04–0.53) and needs a variance-weighted transport space (PCA of log-normalized counts, 10–20 components). FA remains acceptable for the DSM drift of the v56 analyses.
6. **New benchmarks:** a generator calibrated to six real datasets, an independent simulator (dyngen), held-out seeds and pre-registered gates throughout.
7. **Not carried over:** the scJDO v1 analyses of v56 (iPSC reprogramming operator handoffs, hematopoiesis, Dynamo/SpliceJAC concordance, synthetic benchmarks of the v1 machinery, constructed obstruction, bifurcation-saddle limit and K562 Schrödinger-bridge feasibility) are not repeated here. They were obtained with a different operator (a DSM drift fitted on pseudotime), remain available in version 56 of this preprint, and are not re-evaluated in this version.

## Introduction

Cellular identity emerges through regulatory programs that evolve along developmental trajectories. Single-cell RNA sequencing (scRNA-seq) yields high-resolution snapshots of these processes, but a central question remains: where along a trajectory is the system most sensitive to perturbation? Identifying transient windows of instability matters for directed differentiation, for timing interventions and for characterizing the molecular correlates of fate decisions, with causal attribution requiring perturbational follow-up.

Two families of methods address this question. The first reconstructs a scalar landscape or potential: landscape–flux theory decomposes a non-equilibrium drift into a gradient of a potential and a curl flux [5–7], and single-cell methods estimate potentials or energies from snapshots or time courses, including scEpath [8], Topographer [9], population balance analysis [10], MuTrans [11], PRESCIENT [12] and landscape reconstructions from time-evolving distributions [13]. ddHodge reconstructs from RNA velocity a potential, its per-point Hessian, curl and divergence [14], and Zhu et al. quantify landscape, curl flux and entropy production from RNA-velocity vector fields [15]. The second family differentiates a velocity field. Dynamo computes per-cell Jacobians of a vector field fitted to splicing or metabolic-labeling velocity [16–18]; spliceJAC computes state-level Jacobian spectra in which positive eigenvalues mark transition states [19]; and Barcenas et al. locate a tipping point where the leading eigenvalue changes along a lineage [20]. That a leading eigenvalue approaching zero marks a critical transition is a classical idea [21] with several single-cell applications [22–24]. In the previous version of this work (scJDO v1) [1], we computed time-indexed Jacobians of a drift learned by diffusion score matching on pseudotime and decomposed them into recurrent operator archetypes.

Three observations motivate the present revision. First, a Jacobian is only as meaningful as the velocity it differentiates. Snapshot methods differentiate a field derived from the density (a score); kinetic methods differentiate a velocity derived from splicing or labeling. These are different operators with different invariances. The Hessian of the log density is symmetric, and only the signs of its eigenvalues are invariant under a change of coordinates (Sylvester’s law of inertia), whereas a flow Jacobian transforms by similarity and its eigenvalues are invariant. A density fixes neither rotation nor the velocity that generated it [10, 25], and its curvature reflects stability only for a system near a detailed-balance steady state. The per-point Hessian in ddHodge is the Hessian of the gradient part of a velocity field and therefore requires velocity [14]; we found no study that compares the curvature of the density itself, as a separate operator, with an independently measured flow operator per cell.

Second, many datasets have sampling times but no reliable kinetics. Optimal transport (OT) between sampling days provides cell-level couplings and velocities without labels [2, 26, 27], and several methods learn continuous dynamics from time courses by OT or related objectives [3, 4, 12, 28–31]. TIGON and DiffusionOT differentiate the learned velocity, but use the Jacobian to infer gene regulatory networks [3, 4]. Score-based methods separate force from intrinsic noise in time-resolved cross-sections, with force equal to the probability-flow velocity plus a diffusion term times the score [32, 33], and learned scores give local entropy production rates [34]. We found no study that tests whether per-cell eigen-descriptors of a drift fitted to an OT-derived velocity carry usable local information, checked against lineage.

Third, claims about local operators need independent truth. Snapshot non-identifiability is established [10, 25, 35], and benchmarking of Jacobian methods on ground truth shows that eigenvalue-level behavior can be captured while matrix entries are not [36]. We therefore evaluated every operator read-out against held-out seeds of a generator calibrated to real data, an independent simulator and lineage barcodes, pre-registered every gated test, label post hoc analyses as such, and report failures alongside successes.

Here we present scJDO2. To our knowledge, it is the first framework that computes, per cell, both a symmetric score operator (the Hessian of a score-matched log density, compared across representations only through its inertia) and a flow operator (the Jacobian of a drift fitted to a measured velocity, including a label-free entropic-OT velocity), places them in one operator space of magnitude-free descriptors with archetypes, and tests against independent truth, with pre-registered gates, what such operators can and cannot identify. Each ingredient on its own — score matching, OT velocities, velocity-field Jacobians and their eigenvalues, the landscape–flux decomposition and operator archetypes — has precedent, which we cite. We do not claim the first landscape, the first per-cell curvature or Jacobian, the first Jacobian of an OT-learned drift, the leading eigenvalue as a new principle, or a new non-equilibrium measure.

The main findings are that the flow operator estimated from sampling times alone ranks branching cells on unseen data without tuning, has a velocity and drift that predict clonal displacement and a held-out day, orders cell types consistently and scales to a full dataset; that it does not recover the values of the true Jacobian and depends on the transport metric; and that the score operator, tested against knockdowns and clonal fate, did not behave as a measure of biological stability.

## Results

### 1. Two operators, one operator space

Figure 1 summarizes the framework. Which local operator can be computed depends on the time information in the data (Fig. 1).

**Figure 1.**
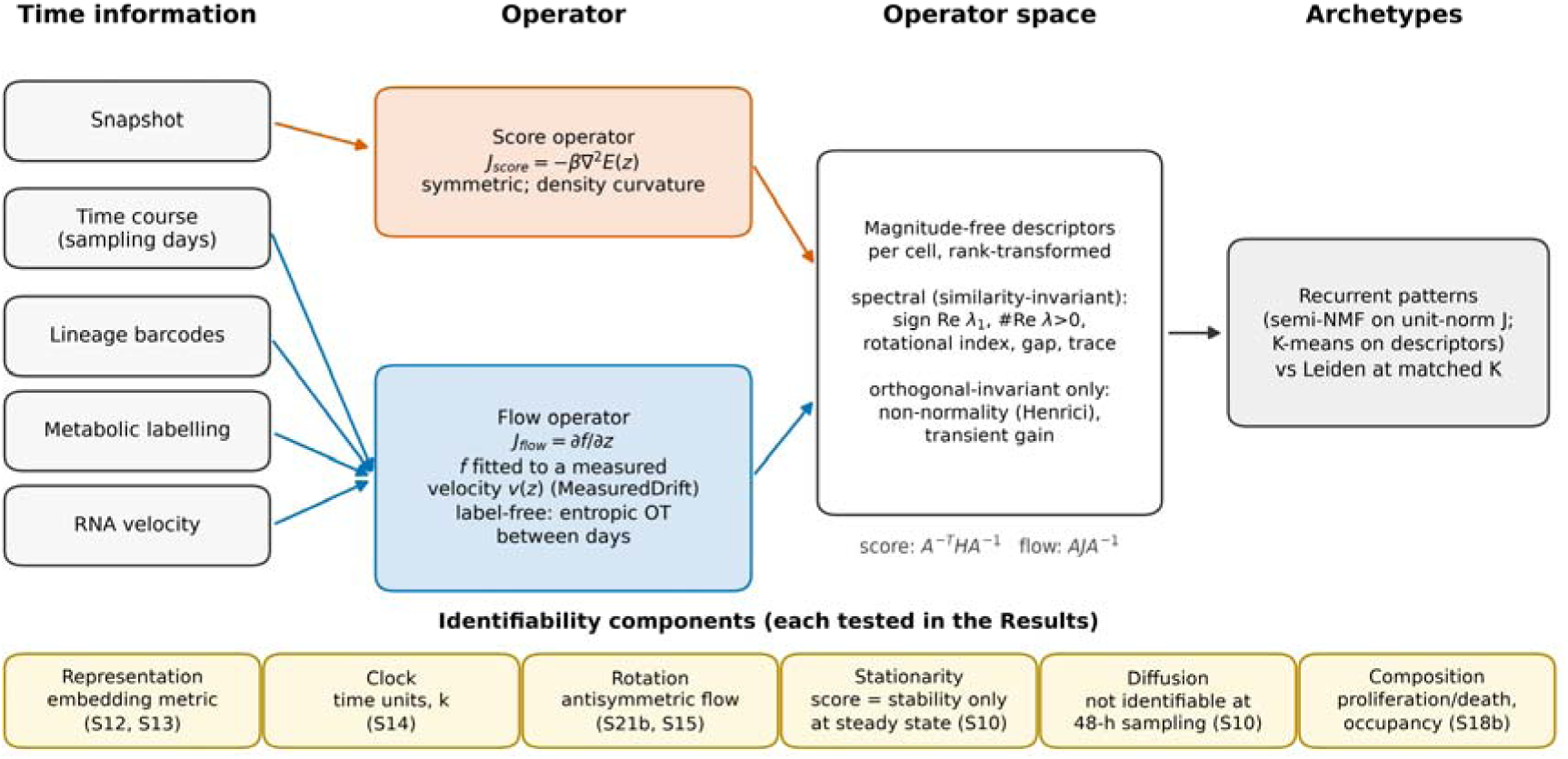
Two operators, one operator space. Schematic (no data). Time information determines which local operator can be estimated. A snapshot supports only the **score operator** *J_score = −β□²E(z)*, the Hessian of a neural energy fitted by score matching; it is symmetric and describes density curvature. A **measured** velocity (entropic optimal transport between sampling days, lineage displacement, metabolic labeling or RNA velocity) supports the **flow operator** *J_flow = ∂f/∂z* of a smooth drift *f* fitted to that velocity (MeasuredDrift). Per cell, magnitude-free descriptors, rank-transformed, define an operator space whose recurrent patterns are archetypes (compared with Leiden clustering at matched K). Under a change of coordinates *A* the score operator transforms as *A⁻^T^HA⁻¹* (congruence) and the flow operator as *AJA⁻¹* (similarity). Spectral descriptors (sign of Re λ□, number of eigenvalues with positive real part, rotational index, spectral gap, trace) are invariant for the flow operator under any invertible *A*; non-normality (Henrici) and transient gain only under orthogonal *A*; for the score operator only the inertia is invariant, and only inertia descriptors are compared between the two operators. Different embeddings (PCA, scVI, FA) are not related by such maps. Bottom: the six identifiability components and the tasks in which each is tested.

#### Score operator

From a snapshot, a neural energy *E(z)* is fitted by score matching [37, 38] to cells in a low-dimensional embedding *z*, and *J_score(z) = −β□²E(z)* is evaluated by automatic differentiation. *J_score* is symmetric. It describes the local curvature of the cell-state density — stiff directions and soft or saddle directions — and coincides with the Jacobian of the dynamics only for a gradient system at a detailed-balance steady state [5, 7]. Out of equilibrium it describes density geometry.

#### Flow operator

When a velocity *v(z)* is measured — by OT between sampling days, lineage displacement, metabolic labeling or RNA velocity — a smooth drift *f* is fitted to *v* (MeasuredDrift), and *J_flow(z) = ∂f/∂z* is computed by automatic differentiation at every cell. *J_flow* is generally non-symmetric and can carry rotation. With splicing or labeling velocity it is a Dynamo-style Jacobian [16]; here we focus on the velocity obtained from sampling times alone (Results 3).

#### Operator space

Under an invertible change of embedding coordinates *A*, the score operator transforms by congruence (*A⁻^T^HA⁻¹*) and the flow operator by similarity (*AJA⁻¹*). Flow-operator eigenvalues are therefore coordinate-invariant, whereas for the score operator only the inertia (the numbers of positive, negative and zero eigenvalues) is. Per cell, we compute eight magnitude-free descriptors of each operator, rank-transformed within each dataset (Methods); their invariance differs. Six are functions of the eigenvalues alone — sign of Re λ□, number of eigenvalues with positive real part, Re λ□/max|λ|, rotational index max|Im λ|/max|λ|, spectral gap/max|λ| and trace/Σ|λ| — and are therefore invariant for the flow operator under any invertible linear change of coordinates. Two — Henrici non-normality and transient gain — depend on the eigenvectors and are invariant only under orthogonal changes of coordinates; under a non-orthogonal map such as whitening or a shear they change (a normal operator can acquire non-zero non-normality). For the score operator only the inertia (sign of the leading eigenvalue and number of positive eigenvalues) is invariant under congruence, and only these inertia descriptors are compared between the two operators. None of these invariances relates different embeddings: PCA, scVI and Factor Analysis coordinates, or PCA spaces with different numbers of components, are not linear (let alone orthogonal) transforms of one another. Descriptors are therefore compared only within one embedding, and the dependence of the operator on the embedding is tested empirically (Results 3). These descriptors define an operator space in which archetypes — the recurrent operator patterns introduced in scJDO v1 [1] — are fitted (Results 7). Within one fixed embedding we also rank cells by the leading eigenvalue λ□ of either operator (Results 3, 4 and 6).

#### Identifiability

Six components limit what can be identified from data: representation (the metric of the embedding), clock (time units), rotation (antisymmetric flow, invisible to a density), stationarity (the score describes stability only at steady state), diffusion, and composition (proliferation and death). Each has precedent in theory [10, 25, 35, 36]. Representation is tested in Results 3, rotation in Results 4, stationarity in Results 3 and 6 and diffusion in Supplementary Fig. S1. Clock and composition were not tested: we use only clock-free (sign- and rank-based) descriptors, and we assume that proliferation and death do not dominate the transport between sampling times (balanced OT).

##### Box 1 What scJDO2 estimates, and what it does not

*Supported under the tested conditions:* a ranking of cells at a fate bifurcation above committed cells by λ□ of the OT flow operator, in a variance-weighted transport space (PCA of log-normalized counts, 10–20 components), without tuning (Results 3, 4); agreement of the drift’s predicted displacement with clonal displacement beyond population movement (a test of the velocity, not of λ□), per-cell λ□ stable across fit seeds and estimators on the same velocity, and cell-type ordering consistent across velocity sources and dataset size (Results 5); a rotational signal on two of three simulated cycles at eight time points (Results 4); a flow archetype that isolates bifurcation cells (Results 7).

*Not supported or not shown:* the values of the true Jacobian (flow or score); per-cell reproducibility of flow operators across independent velocity sources on real data; the timing of a critical transition (no confirmatory test could be constructed); diffusion; the representation-independence of the flow operator (it inverts on Factor Analysis); the score operator as a predictor of knockdown responses or clonal fate; archetypes as a validated regime classifier.

### 2. A benchmark calibrated to real data

We measured 20 statistics (32 sub-statistics) on six real datasets (scNT-seq hematopoiesis [16] and neurons [39], LARRY [40], Setty CD34⁺ [41], pancreas [42] and dentate gyrus [43]): expression level and sparsity, dimensionality, count noise, gene programs, splicing-fit difficulty, turnover and lineage structure (Fig. 2a). Earlier generator versions were unrealistic; for example, the median zero fraction was 36–37 % in v1 and v2 data against 84–96 % in real data. The calibrated generator (v3) matches expression level, zeros, noise, barcode rate, clone size and labeling signal (Fig. 2a,b). Against 13 calibration targets fixed in advance it met 6 of 13 on each seed, failing the pre-registered realism gate; the targets missed on all three seeds were genes detected, the fraction of clones spanning several time points, the unspliced fraction, the share of steady genes and the scVelo recovered fraction. Its overall match rate was 0.39 (v1 0.35, v2 0.22; Fig. 2c). We accepted v3 with these misses documented, and state them next to every splicing- or lineage-based number. An independent simulator, dyngen [44], matched the same statistics at 0.51 (range 0.44–0.60). The v3 presets include a pitchfork bifurcation with a controlled critical point (transition), a no-fork null, batch effects, stress, a stationary cycle and time-course sampling (days 2, 4 and 6) with lineage barcodes and labeling.

**Figure 2.**
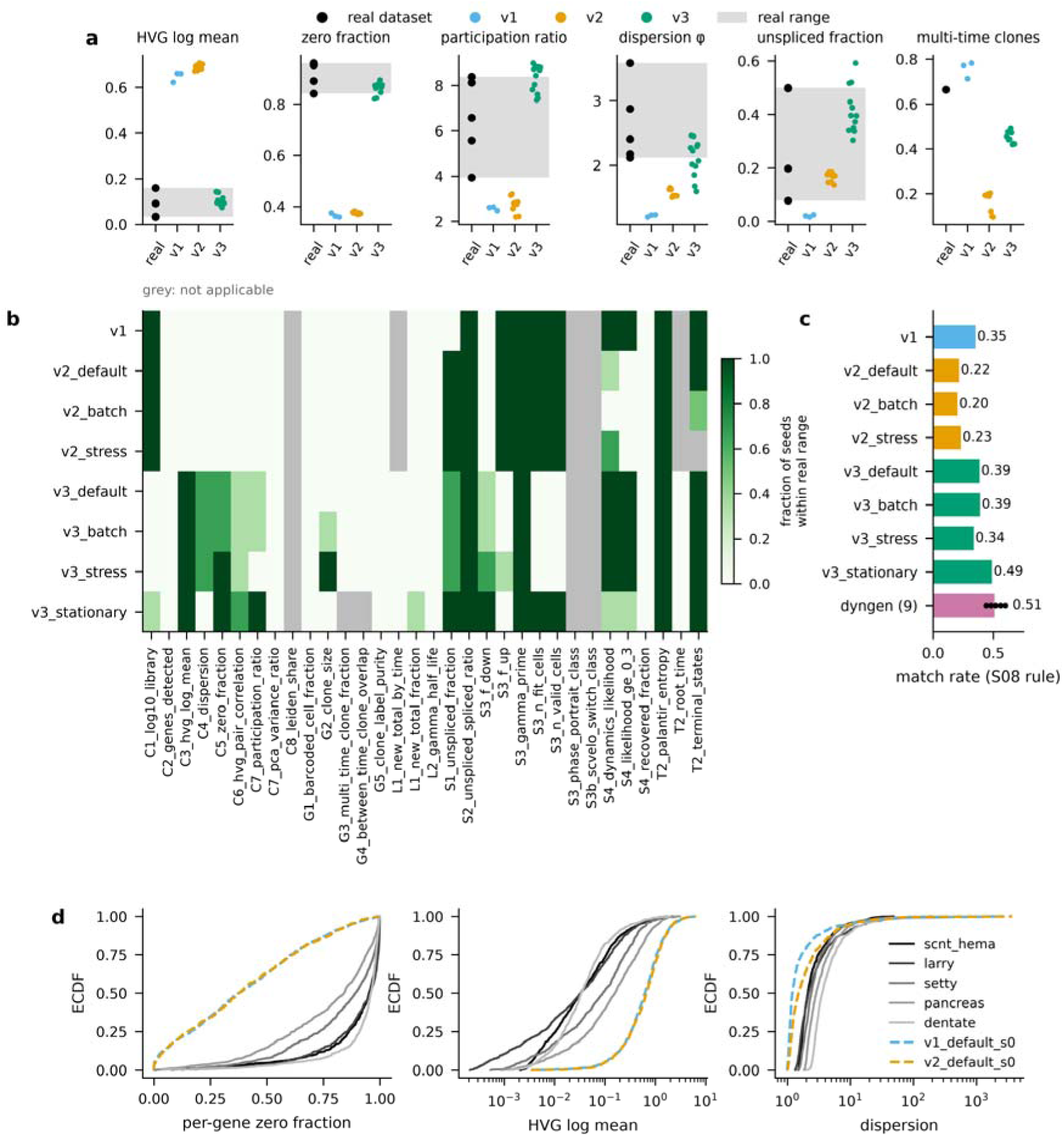
A benchmark calibrated to six real datasets (S08, S09, S19). **a**, Six of the 20 S08 statistics (IDs C1–C8, G1–G5, L1–L2, S1–S4, T2; 32 sub-statistics). Black dots: value (or median) in each real dataset (scNT-seq hematopoiesis and neurons, LARRY, Setty CD34⁺, pancreas, dentate gyrus; not every statistic applies to every dataset); gray band: real min–max range. Colored dots: one per synthetic dataset (v1: 3 seeds of the default preset; v2 and v3: 3 seeds × each preset). Zero fraction: real 0.84–0.96 vs v1 median 0.36, v2 0.37, v3 0.87. **b**, Whether the synthetic median lies in the real range, per statistic and preset (fraction of three seeds); gray, not applicable. **c**, Fraction of statistics matched under the S08 rule (distance ≤ max(real spread, fixed tolerance) for every applicable real reference): v1 0.35, v2_default 0.22, v3_default 0.39, v3_batch 0.39, v3_stress 0.34, v3_stationary 0.49; dyngen (S19, nine datasets, bifurcating/cycle/linear × 3 seeds, 25 applicable statistics) 0.51 (range 0.44–0.60). Against the 13 pre-fixed S09 calibration targets, v3_default met 6 of 13 on each seed (realism gate failed); targets unmet on all three seeds: C2_genes_detected, G3_multi_time_clone_fraction, S1_unspliced_fraction, S3_steady_share, S4_recovered_fraction. The authors accepted v3 with these misses documented (S09 decision). **d**, Empirical CDFs of three saved S08 distributions for five real datasets and v1/v2 default seed 0 (v3 distributions were not saved). Descriptive comparison; the S09 13-target count was pre-registered, the S08/S19 matrices are report only.

### 3. The flow operator from a time course alone ranks branching cells, untuned

#### Label-free estimate

Entropic OT (Sinkhorn; ε = 0.05 × median squared distance) between consecutive sampling days in a PCA of log-normalized counts gives each cell a barycentric velocity; a drift is fitted to these velocities and differentiated (Methods).

#### Held-out test

The pre-registered test (Fig. 3a) compared two configurations on held-out seeds never used for selection: a reference configuration fixed a priori (the drift and transport defaults of an earlier task, PCA with k = 15) and a configuration tuned one factor at a time on two development seeds. On the transition preset, λ□ of the reference flow operator separated cells at the true saddle from committed cells with AUC 0.89–0.97 across k = 10–20 principal components, and 0.97–1.00 on the default, batch and stress presets. The tuned configuration did worse on the same held-out seeds (0.42–0.95). All four pre-registered gates, which applied to the tuned configuration and its development checks, failed. The rule for choosing k was amended after development AUCs had been seen; test seeds were untouched (Methods). The reading is that tuning on two development seeds overfits and the a-priori settings are robust.

**Figure 3.**
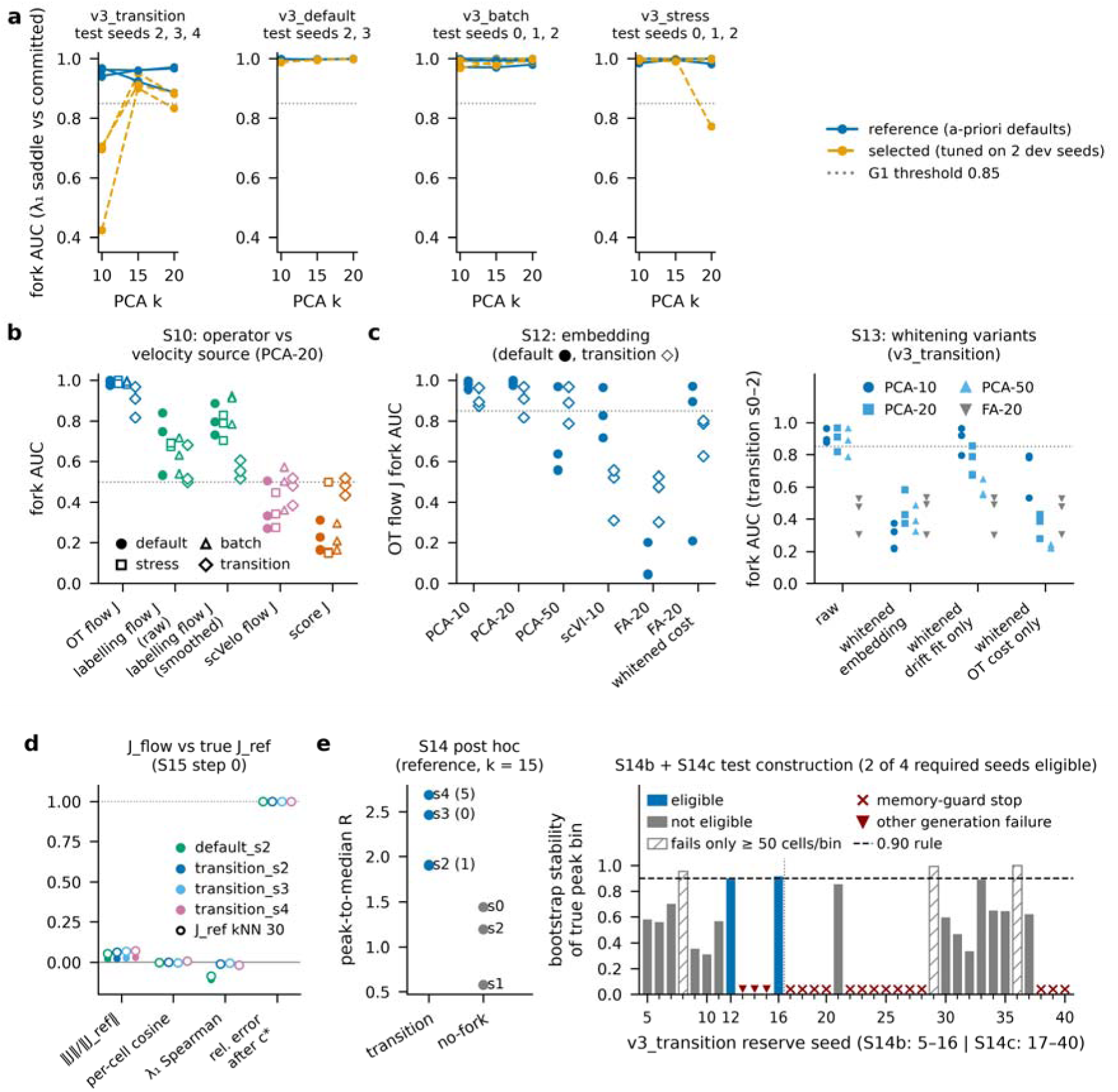
The flow operator from a time course alone ranks branching cells, untuned (S10, S12–S15, S14b, S14c). Flow J = Jacobian of MeasuredDrift fitted to entropic-OT barycentric velocities between sampling days (days 2/4/6) in PCA of log-normalized counts. Fork AUC = AUC of the leading eigenvalue λ□ for true saddle vs committed cells (truth from generator state, used for scoring only). **a**, S14 held-out test (pre-registered; test seeds never used for selection): 11 test datasets (v3_transition s2–s4, v3_default s2–s3, v3_batch s0–s2, v3_stress s0–s2) at k = 10, 15, 20. Reference (a-priori) configuration: AUC 0.89–0.97 on v3_transition and 0.97–1.00 on the other presets. Tuned configuration (one factor at a time on two development seeds): 0.42–0.95 on v3_transition (gate G1, ≥ 0.85 on every transition seed at k* = 15 and neighbors, failed; minimum 0.42). **b**, S10 (pre-registered gate on labeling flow J; other operators report only): fork AUC per operator on v3 presets (PCA-20; 3 seeds each; filled = v3_default). v3_default: OT flow J 0.97–1.00, labeling flow J 0.53–0.89 (raw and smoothed variants), scVelo flow J 0.27–0.51, score J 0.16–0.31. **c**, Left, S12 Step 0 (6 datasets: v3_default and v3_transition s0–s2): OT flow J fork AUC by embedding — PCA-10 0.88–1.00, PCA-20 0.82–1.00, PCA-50 0.56–0.97, scVI-10 0.31–0.96, FA-20 0.04–0.53, FA-20 with noise-whitened cost 0.21–0.97. Right, S13 (pre-registered; gate failed) on v3_transition s0–s2: whitening the embedding (PCA-20 0.37–0.58), whitening only the drift fit (PCA-10 0.79–0.96), whitening only the OT cost (PCA-10 0.53–0.79, PCA-20 0.28–0.43). **d**, Flow J vs the Jacobian of the generator’s true expected velocity in the same PCA-15 coordinates (J_ref; S15 Step 0; 4 test datasets × J_ref neighborhood 15/30): norm ratio 0.02–0.07, median per-cell cosine −0.002 to 0.008, per-cell λ□ Spearman −0.107 to −0.005 across all four datasets (−0.018 to −0.005 on the three transition seeds), relative error after optimal scalar c* 1.00. The reference was verified by finite differences of the true expected velocity (median per-cell cosine ≈ 1.000). Across S14 test rows the median relative Frobenius error is 1.000–1.005. **e**, Left, *post hoc*: peak-to-median ratio R of the 12-bin day-weighted strict-pseudotime λ□ profile (reference, k = 15) on the three transition test seeds (1.91–2.69; |detected − true bin| 1, 0, 5) and three no-fork seeds (0.58–1.44). Right, S14b and S14c construction (pre-registered): bootstrap stability of the true critical peak bin per reserve seed (dashed: 0.90 rule). S14b: 2 of 9 assessed seeds eligible (seeds 12, 16); seeds 13, 14, 15 could not be generated. S14c (seeds 17–40): 14 seeds stopped at the generation memory guard; none of the 10 assessed was eligible; seeds with stable interior peaks failed only the ≥ 50-cell bin rule (minimum bins 42–46 cells). Four eligible seeds were required; no model was fitted, and the confirmatory transition test was not constructed.

#### Velocity sources

On the default preset (PCA-20; Fig. 3b), the OT flow operator reached fork AUC 0.97–1.00, whereas flow operators from labeling velocity (0.53–0.89) or scVelo velocity (0.27–0.51) were weaker. The score operator ranked committed cells above saddle cells (0.16–0.31), as expected for a system far from steady state (Results 6).

#### Representation

The result depends on the transport metric (Fig. 3c; Supplementary Fig. S3). Fork AUC held in PCA-10 (0.88–1.00) and PCA-20 (0.82–1.00) and degraded in PCA-50 (0.56–0.97), scVI-10 [45] (0.31–0.96) and Factor Analysis (FA-20; 0.04–0.53, inverted), although OT velocities agree between PCA and FA in gene space (0.52–0.56). In a pre-registered follow-up (gate failed), whitening the embedding destroyed the signal (PCA-20 0.37–0.58), whitening only the drift fit preserved it (PCA-10 0.79–0.96), and whitening only the OT cost degraded it (PCA-10 0.53–0.79; PCA-20 0.28–0.43). The signal is carried by the variance-weighted OT metric, which equal-variance representations (FA, whitening) remove. This corrects v56’s recommendation of FA as the default for operator inference.

#### What the operator is not

We compared the flow operator with the Jacobian of the generator’s true expected velocity in the same coordinates (*J_ref*; Fig. 3d). The median per-cell relative error is ≈ 1.0 on every held-out dataset (1.000–1.005). The flow operator’s norm is 2–7 % of the reference’s, optimal rescaling does not reduce the error, the median per-cell cosine is ≈ 0 (−0.002 to 0.008) and the per-cell λ□ Spearman correlation is −0.107 to −0.005 across all four datasets (−0.018 to −0.005 on the three transition seeds; −0.107 and −0.086 on v3_default s2). The reference was verified by an independent route — finite differences of the true expected velocity in the same coordinates, with median per-cell cosine ≈ 1.000 against *J_ref* — so the discrepancy lies in the estimate, not the reference. The flow operator therefore does not recover the Jacobian’s values; its λ□ ranks branching cells empirically. Consistent with eigenvalue-level benchmarks of Jacobian methods [36], all operator-space descriptors in this paper are sign- or rank-based.

#### Transition detection is not shown

Post hoc, on the three held-out transition seeds and three no-fork seeds, the peak-to-median ratio R of binned λ□ along strict pseudotime separated all transitions (1.91–2.69) from all no-forks (0.58–1.44), with timing offsets of 0, 1 and 5 bins (Fig. 3e, left). Because this comparison was chosen after the results, we treat it as exploratory. Two pre-registered confirmatory tests were designed to require four transition datasets that were eligible before any model was fitted: strict-pseudotime provenance complete, at least 50 cells in every one of 12 bins, a true critical peak in an interior bin, and the same peak bin in at least 90 % of 200 bootstrap resamples. In the first attempt, only 2 of 9 assessed seeds were eligible; the true critical peak was usually flat on the 12-bin scale, and three further seeds failed in the generator (Fig. 3e, right). In the second attempt, an extended reserve of 24 seeds yielded no further eligible dataset: 14 seeds stopped at the memory guard during generation and none of the 10 assessed was eligible. Across both attempts, the three seeds with stable interior peaks (bootstrap stability 0.955–1.000) failed only the bin-occupancy rule, with minimum bins of 42–46 cells set by about 180 day-2 cells. The occupancy rule thus excluded exactly the well-ordered datasets, a design error identified at review. Under the pre-registered stopping rule, no model was fitted in either attempt. Transition detection is therefore not shown, and we make no claim about the timing of critical transitions.

### 4. An independent simulator

On dyngen data (5,000 cells, 1,000 genes, eight synchronized sampling times; truth used for scoring only), with the same untuned settings (Fig. 4):

- **Bifurcations.** Flow-operator fork AUC was 1.00, 0.92–0.99 and 0.31–0.38 (inverted) on the three datasets, consistent across three drift-fit seeds; the pre-registered gate (≥ 0.85 on every dataset) failed. The score operator reached 0.98, 0.58 and 0.85 (Fig. 4a). The inverted dataset has not yet been diagnosed.
- **Cycles.** The flow operator’s rotational index was higher on cycles than on linear trajectories for two of three cycles (0.19–0.40 vs 0.07–0.09) but not for the third (0.02 vs 0.08–0.10); the gate failed (Fig. 4b). The score operator is symmetric and has no rotation.
- **Linear trajectories.** No cell had Re λ□ > 0, under any drift-fit seed (Fig. 4c).

**Figure 4.**
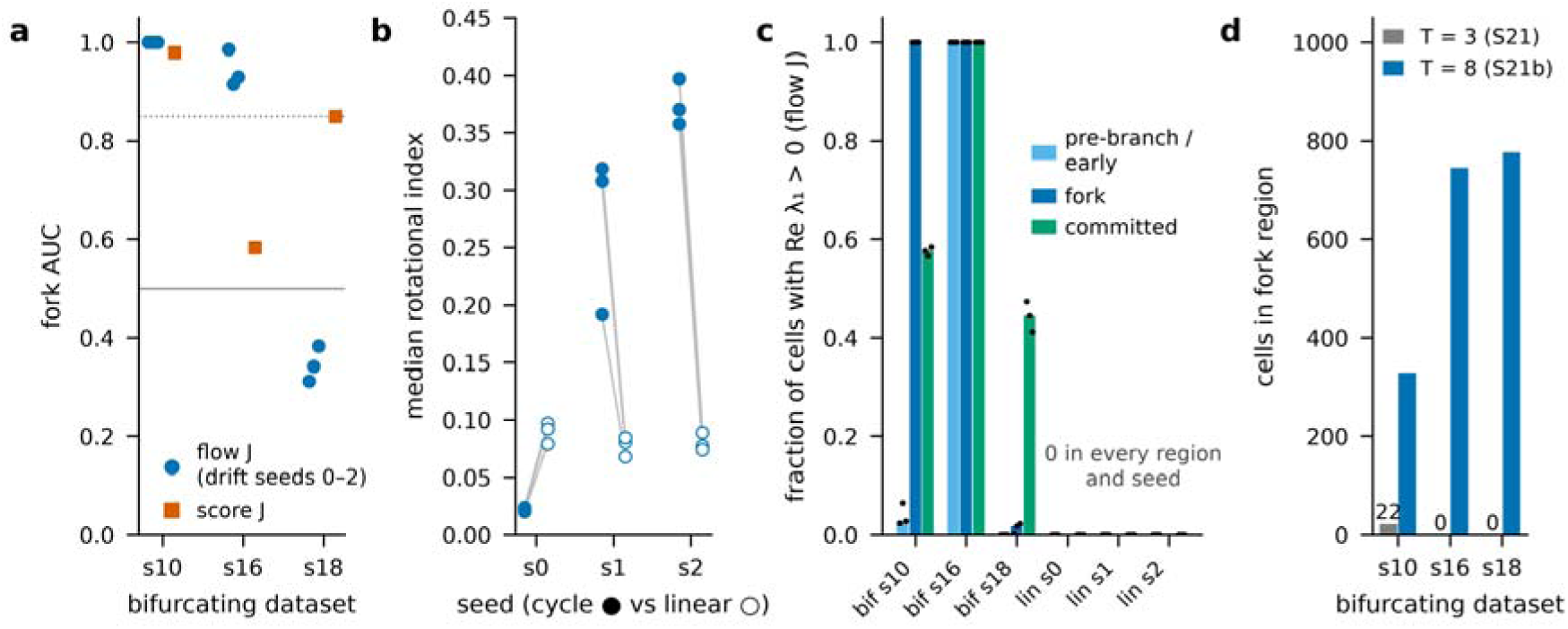
An independent simulator (dyngen; S21b, pre-registered). Nine dyngen datasets (bifurcating seeds 10/16/18, cycle s0–s2, linear s0–s2; 5,000 cells, 1,000 genes), re-sampled from the saved S21 simulations at eight synchronized time points; same untuned settings as S14’s reference flow J (PCA-15, entropic OT, MeasuredDrift drift seeds 0–2) and S10 score J; truth read only after all fits. **a**, Fork AUC (λ□, fork vs committed cells): flow J 1.00, 0.92–0.99 and 0.31–0.38 (inverted) on s10, s16, s18; score J 0.98, 0.58, 0.85. Gate F1 (≥ 0.85) holds on 2 of 3 datasets (gate failed). **b**, Median rotational index max|Im λ|/max|λ| of flow J, cycle (filled) vs linear (open) per dataset seed and drift seed (lines pair the same drift seed): cycle s1 0.19–0.32 and s2 0.36–0.40 vs linear 0.07–0.09; cycle s0 0.02 vs linear s0 0.08–0.10 (gate F2 failed: 2 of 3 cycles pass on every seed). Score J is symmetric (rotational index 0.00). **c**, Fraction of cells with Re λ□ > 0 (flow J) by truth region (report only): on linear backbones the fraction is 0 in the early and committed regions for every seed, and 0 over all 5,000 cells (maximum 0.0000 over 9 dataset × seed rows). **d**, Cells in the fork region at three time points (S21) vs eight (S21b): s10 22 vs 329, s16 0 vs 746, s18 0 vs 776. The inverted dataset (s18) has not yet been diagnosed.

Three sampling times were too coarse (Fig. 4d; Supplementary Fig. S2). They placed 22, 0 and 0 cells at the branch point (against 329, 746 and 776 at eight times), so the fork test was evaluable on one dataset only; the rotational index of cycles stayed at the level of linear trajectories; and every early (pre-branch) cell had Re λ□ > 0. With eight time points this fraction fell to 0.00–0.06 on two of three datasets (1.00 on the third), and we withdrew the three-time-point reading as a sampling artifact.

### 5. Real data: lineage agreement, held-out prediction and scale

*LARRY* (in vitro hematopoiesis, days 2, 4 and 6 [40]; dense analyses use a fixed sample of 4,000 cells per day; barcodes used for scoring only; Fig. 5a,b):

- **Velocity.** OT and clonal-displacement velocities agree (median cosine 0.69 over 1,321 cells with a lineage velocity).
- **Lineage.** Integrating the reference drift over each 48-h interval from a barcoded cell predicts its displacement to the centroid of the same clone’s cells at the next sampling day (pre-registered statistic: Spearman 0.56, pooled over the 15 PCA coordinates of 554 cell–clone pairs; maximum over five permutation nulls 0.15). Post hoc, 1,000 permutations give a one-sided p < 0.001 (null maximum 0.15). Because pooled coordinates also reward moving with the population, we added controls: a population mean-shift predictor reaches 0.19; after centring every coordinate within each interval, the drift still agrees with clonal displacement (0.50; null maximum 0.04); the OT displacement without a drift reaches 0.54 (0.48 centred); and displacement magnitudes agree weakly (0.24). λ□ is unrelated to displacement magnitude (−0.04). Lineage therefore validates the OT velocity and drift cell by cell beyond population movement; it does not test λ□ or branching (script scripts/s14_lineage_addendum.py).
- **Prediction.** Integrating the fitted drift from day 4 predicts the held-out day-6 distribution better than mean-displacement or no-movement baselines (energy distance 0.69 vs 0.87 and 0.88; post hoc 95 % bootstrap intervals 0.66–0.73, 0.85–0.89 and 0.83–0.94, with the flow best in all 200 resamples). Transport and drift were fitted on days 2 and 4 only (Methods). The tuned configuration did not (0.94), and the gate, which applied to it, failed.
- **Stability.** Fits are stable across drift-fit seeds and estimators (λ□ Spearman 0.92–0.95, including Dynamo’s SparseVFC on the same velocity [16]), and both the OT and the lineage flow operators rank undifferentiated cells above neutrophils and monocytes.
- **Per cell.** The flow operators from OT and from lineage velocity agree only at null level (λ□ Spearman 0.27, null maximum 0.27; leading-eigenvector |cos| 0.29, null maximum 0.32; Supplementary Fig. S4). Coarse, cell-type-level structure is reproducible; per-cell operators across independent velocity sources are not. This pre-registered real-data test met 6 of 14 criteria.

**Figure 5.**
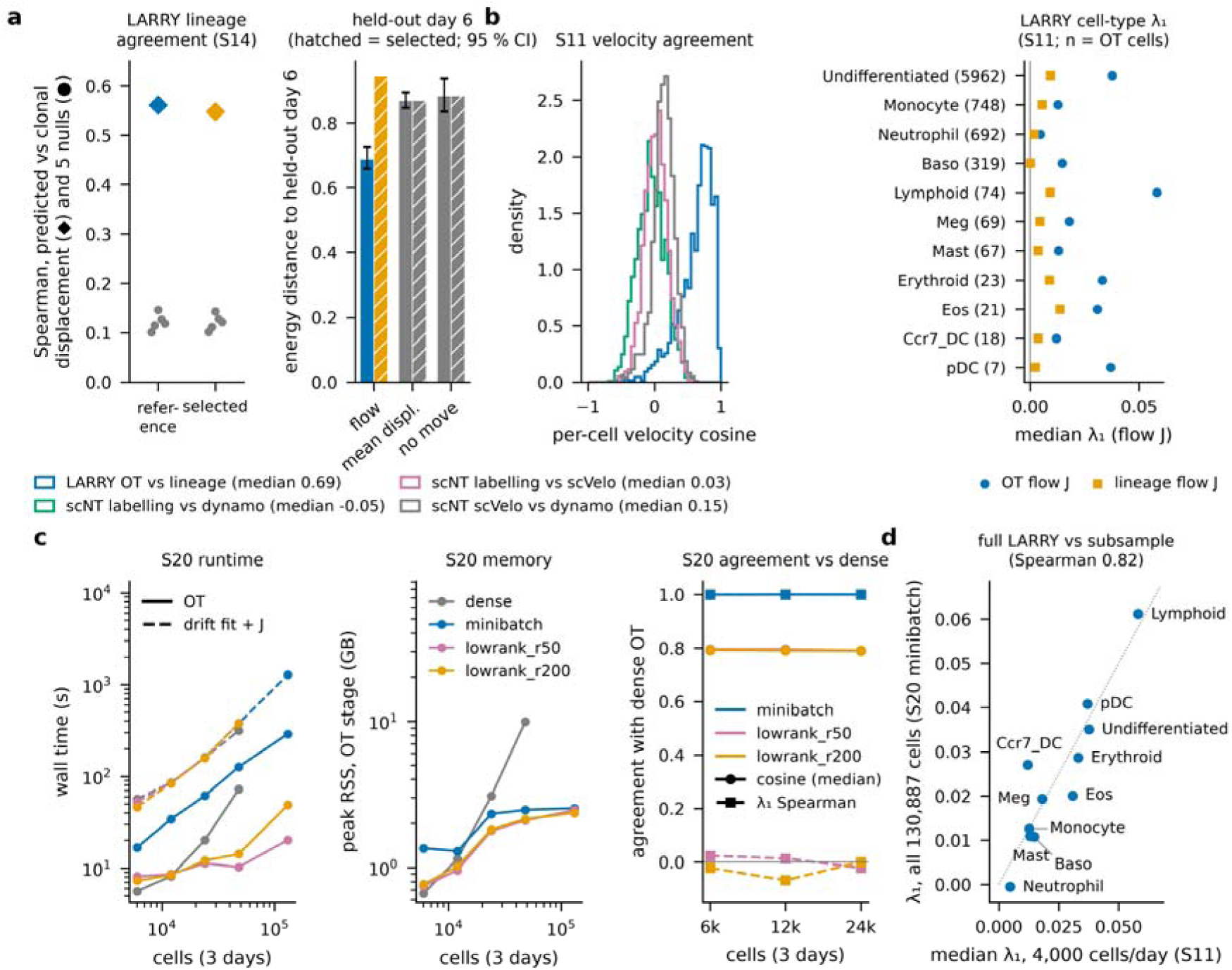
Real data: lineage agreement, held-out prediction and scale (S14, S11, S20). LARRY in vitro hematopoiesis (GSE140802; days 2/4/6); dense analyses use a fixed sample of 4,000 cells per day (default_rng(0)); barcodes used for scoring only. **a**, S14 held-out test (pre-registered; gate G3 applies to the tuned configuration and failed). Left, Spearman, pooled over PCA coordinates of 554 barcoded cell–clone pairs, between the displacement predicted by integrating the drift over the 48-h interval and the observed displacement to the clone’s next-day centroid (□), vs five permutation nulls (●; pre-registered): reference 0.56 (null max 0.15), tuned 0.55 (null max 0.14). Post hoc, reference: 1,000 permutations, one-sided p < 0.001 (null max 0.15); after centring each coordinate within an interval 0.50 (null max 0.04); population mean-shift predictor 0.19; OT displacement without a drift 0.54; λ□ vs displacement magnitude −0.04. This tests the velocity and drift, not λ□. Right, energy distance between the day-6 distribution predicted by integrating the fitted drift from day 4 and the held-out day 6: reference flow 0.69, tuned flow 0.94; mean-displacement baseline 0.87, no-movement baseline 0.88. Error bars (reference, post hoc): 95 % bootstrap intervals over cells (200 resamples): flow 0.66–0.73, mean displacement 0.85–0.89, no movement 0.83–0.94. Transport and drift were fitted on days 2 and 4 only; the PCA on all three days. **b**, S11 (pre-registered criteria; 6 of 14 met). Left, per-cell cosine between velocity sources: LARRY OT vs clonal displacement (n = 1,321 cells with a lineage velocity; median 0.69); scNT-seq hematopoiesis labeling vs Dynamo −0.05, labeling vs scVelo 0.03, scVelo vs Dynamo 0.15 (n = 1,947). Right, LARRY cell-type median λ□ of the OT and lineage flow J (PCA-30); Undifferentiated > Neutrophil and Monocyte for both. Per-cell OT vs lineage λ□ Spearman 0.27 (null level); OT flow J across fit seeds 1–2 and Dynamo SparseVFC 0.92–0.95. **c**, S20 (pre-registered gates G1, G2 passed): wall time (OT solid, drift fit + Jacobians dashed) and OT-stage peak memory vs total cells (2,000–16,000 per day and all 130,887 cells) for dense, mini-batch (4,000-cell batches) and low-rank (moscot rank 50/200) OT; dense OT on the full data was not run (cost matrix ≈ 21 GB). Full LARRY with mini-batch OT: OT 291 s (2.54 GB), drift fit + Jacobians 1,278 s (1.31 GB). Right, agreement with dense OT: mini-batch median velocity cosine 0.9998 and λ□ Spearman 0.9999 at 8,000 cells/day (the 4,000/day gate row is trivial: one batch = whole day); low-rank cosine 0.79–0.80, |λ□ Spearman| ≤ 0.07. **d**, Cell-type λ□ on all cells (cell-count-weighted mean of per-day medians, 11 cell types) vs the 4,000-per-day S11 result: Spearman 0.82.

#### *Scale* (Fig. 5c,d)

Mini-batch OT (4,000-cell batches, each cell in ten batches) reproduces dense OT at 8,000 cells per day (median velocity cosine 0.9998; λ□ Spearman 0.9999). It ran on all 130,887 LARRY cells in 291 s (2.5 GB), plus drift fitting and Jacobians in 21 min (1.3 GB); dense OT on the full data was not run (cost matrix ≈ 21 GB). Cell-type λ□ on the full data agrees with the 4,000-per-day result (Spearman 0.82 over 11 cell types). Low-rank OT (moscot, rank 50 or 200 [27]) did not preserve the operator (velocity cosine 0.79–0.80; λ□ Spearman ≈ 0).

#### Labeling data

On scNT-seq hematopoiesis [16], labeling, scVelo and Dynamo velocities disagree before any operator is fitted (median per-cell cosine: labeling vs Dynamo −0.05, labeling vs scVelo 0.03, scVelo vs Dynamo 0.15; Fig. 5b). The labeling velocity depends on whether degradation rates are shared across pulses: with Dynamo’s convention, agreement with Dynamo rises from ≈ 0 to 0.41 (exploratory diagnostic; script coord/orient/reviews/labelling_diag_2026-10-19/). Labeling-based flow operators must be shown robust to this choice before they are interpreted.

### 6. The score operator: a density-curvature descriptor, not validated against perturbation or lineage

By construction, the score operator describes the curvature of the cell-state density, and it reflects dynamical stability only near a detailed-balance steady state. Out of equilibrium it failed as expected: on four non-stationary presets it ranked committed cells above saddle cells (AUC 0.16–0.31 on default, 0.15–0.50 on stress, 0.17–0.29 on batch, with AUC < 0.45 on 8 of these 9 seeds, and 0.43–0.52 on transition; Fig. 6a). On dyngen it reached 0.98, 0.58 and 0.85 on the three bifurcations (Results 4). We then tested whether it carries information about biological stability against two independent truths, both pre-registered:

- CRISPRi knockdowns (Replogle K562 and RPE1 [46]).

`o` In K562 (10,691 non-targeting controls; 865 of 1,299 targets with ≥ 100 cells passed a BH-adjusted permutation test), three soft directions of the local score operator captured a median 4.5 % of each knockdown’s centroid shift, against 58.9 % for the top three directions of the global control covariance and a random-subspace expectation of 15 % (Fig. 6b). The pre-registered criterion (≥ 70 % of targets above their random-null 95th percentile) reached 5.3 %.
`o` Re-weighting control states explained a median 31 % of each knockdown’s energy distance (required ≥ 50 %), and basin stiffness did not predict knockdown-induced occupancy change across 59 basins (partial Spearman ρ +0.05, one-sided p 0.65; Fig. 6c). An operator-free Leiden baseline was equally null.
`o` A dose-response confirmation on RPE1 was underpowered (1 qualifying target; ≥ 10 required). In K562 (report only), score-operator capture of weak-knockdown displacements did not exceed a shared-basis null or the covariance baseline (Fig. 6d; Supplementary Table S3).
`o` A post hoc observation — strong knockdowns of transcription machinery move along soft directions of the score operator computed near their destination, but not from random controls — is hypothesis-generating only.
- Clonal fate coherence (LARRY). Clones whose cells sit in stiffer regions of the score operator do not keep their sisters in fewer fates at the next time point. The pooled correlation between clone λ□ and fate entropy was ρ = −0.003; within undifferentiated cells it was −0.10, the wrong sign. Distance to the progenitor centroid predicted fate spread better (|ρ| 0.22 vs 0.10 for λ□), and the partial correlation given local density was −0.08. All three gates failed (Supplementary Table S2; Fig. 6d).

**Figure 6.**
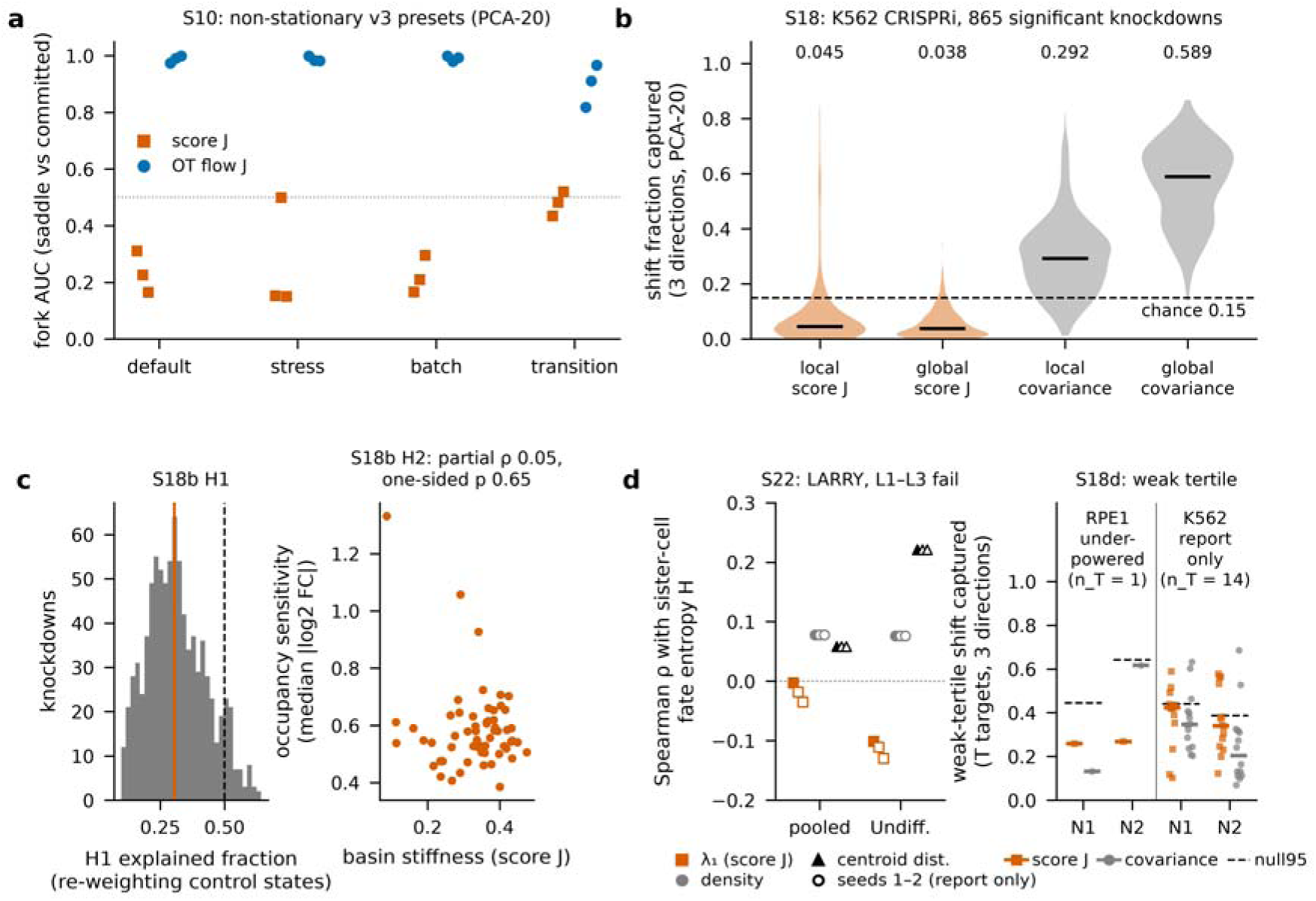
The score operator: a density-curvature descriptor, not validated against perturbation or lineage (S10, S18, S18b, S22, S18d). **a**, S10: fork AUC of score J (−β□²E, energy network fitted by exact-Laplacian score matching) vs OT flow J on four non-stationary v3 presets (PCA-20, 3 seeds each). Score-J AUC on default 0.16–0.31, stress 0.15–0.50 and batch 0.17–0.29 (AUC < 0.45, i.e. committed cells ranked above saddle cells, on 8 of 9 seeds), 0.43–0.52 on transition; OT flow J 0.97–1.00 on default. **b**, S18 (pre-registered; P1 failed): Replogle K562 CRISPRi; all fits on 10,691 non-targeting controls; 865 of 1,299 targets with ≥ 100 cells passed the BH-adjusted permutation test. Fraction of each knockdown’s centroid shift captured by m = 3 directions in control PCA-20: local score-J soft axes 0.045, global score J 0.038, local covariance 0.292, global covariance 0.589 (medians); random-subspace expectation 0.15. P1 (fraction of targets above their random-null 95th percentile ≥ 0.70): 0.053. k = 10/15/30 (report only): local score J 0.099/0.056/0.051. **c**, S18b (pre-registered; both gates failed). Left, H1: fraction of each knockdown’s energy distance explained by re-weighting control states (k = 20): median 0.31 (required ≥ 0.5); 235 of 865 targets (0.27) above the size-matched noise 95th percentile. Right, H2: score-J basin stiffness vs occupancy sensitivity across 59 basins; partial Spearman ρ (ranks residualized on basin size and covariance trace) 0.05, one-sided p (negative) 0.65 (10,000 permutations); ordinary ρ 0.06; operator-free Leiden spread ρ 0.03 (p⁻ 0.60). **d**, S22 (score-J stiffness vs LARRY sister-cell fate coherence; pre-registered, all gates failed) and S18d (RPE1 dose-response confirmation; underpowered). Left, S22 (pre-registered; L1 failed, L2 failed, L3 failed): Spearman ρ between a LARRY clone’s median score-J λ□ on day *t* and the entropy H of its sister cells’ types on day *t* + 2 (filled: seed 0, gated; open: seeds 1–2, report only), next to the local-density and distance-to-Undifferentiated-centroid baselines. Pooled (1,818 clone/time pairs, 67 with tied modal types): λ□ ρ −0.003 (one-sided permutation p 1); Undifferentiated (1,401 pairs): λ□ ρ −0.10 (p 1), centroid distance 0.22, density 0.08; partial ρ of λ□ given density −0.08. λ□ ρ over seeds 0–2 and both cohorts: −0.130 to −0.003. Right, S18d: fraction of the weak-knockdown-tertile centroid shift of each transcription-machinery (T) target captured by three control score-J soft directions (vermillion) or three control-covariance directions (gray); bars, medians over T targets; dashed, 95th percentile of the across-T median under 10,000 covariance-matched random subspaces (shared basis). N1, depth-projected log-normalization; N2, analytic Pearson residuals. RPE1 (pre-registered; underpowered: 1 qualifying T target, ≥ 10 required): score J 0.26/0.27 (N1/N2), covariance 0.13/0.62, null95 0.44/0.64. K562 (report only; 14 T, 38 M targets): score J 0.42/0.34, covariance 0.35/0.20, null95 0.44/0.39; score J above covariance, one-sided paired Wilcoxon p 0.27/0.09; T above mitochondrial/translation (M) targets, one-sided Mann–Whitney p 0.002/0.012. Supplementary Tables S2 and S3.

The score operator therefore describes density curvature, and we do not claim that it measures biological stability. In K562, a proliferating line, the steady-state assumption under which curvature would equal stability need not hold; in LARRY, it is the progenitor’s position rather than its curvature that predicts fate spread.

## 7. Archetypes of dynamics

Operator archetypes were introduced in scJDO v1 for temporal sequences of DSM-drift Jacobians [1]. Here we asked whether archetypes of flow operators recover the true dynamical regimes of the transition preset (Fig. 7). Archetypes were fitted to unit-norm per-cell flow operators by semi-NMF [47, 48] with K chosen by held-out reconstruction (operator-archetype view, K = 8), and by K-means on rank-transformed descriptors (descriptor view, K = 2). They were compared with score-operator archetypes and with Leiden clustering of expression at matched K [49], on the three held-out transition seeds.

- **A flow archetype isolates the saddle.** In the operator-archetype view, one archetype captured saddle cells with recall 0.72–0.79 at 3.4–4.1× their base rate on every test seed (gate A1 passes). The descriptor view had higher recall (0.88–0.95) but failed precision on one seed (1.5×).
- **It does not beat expression clustering overall.** Against all true regimes, normalized mutual information was 0.35–0.39 for flow archetypes, 0.35–0.37 for Leiden at matched K and 0.05–0.31 for score archetypes. Flow beat score on every seed but Leiden on only one (gate A2 fails).
- **Archetypes are not stable across drift-fit seeds.** The adjusted Rand index of assignments between seeds was 0.22–0.44 in the operator view and −0.02 to 0.53 in the descriptor view (gate A3, ≥ 0.5 for every pair, fails).

**Figure 7.**
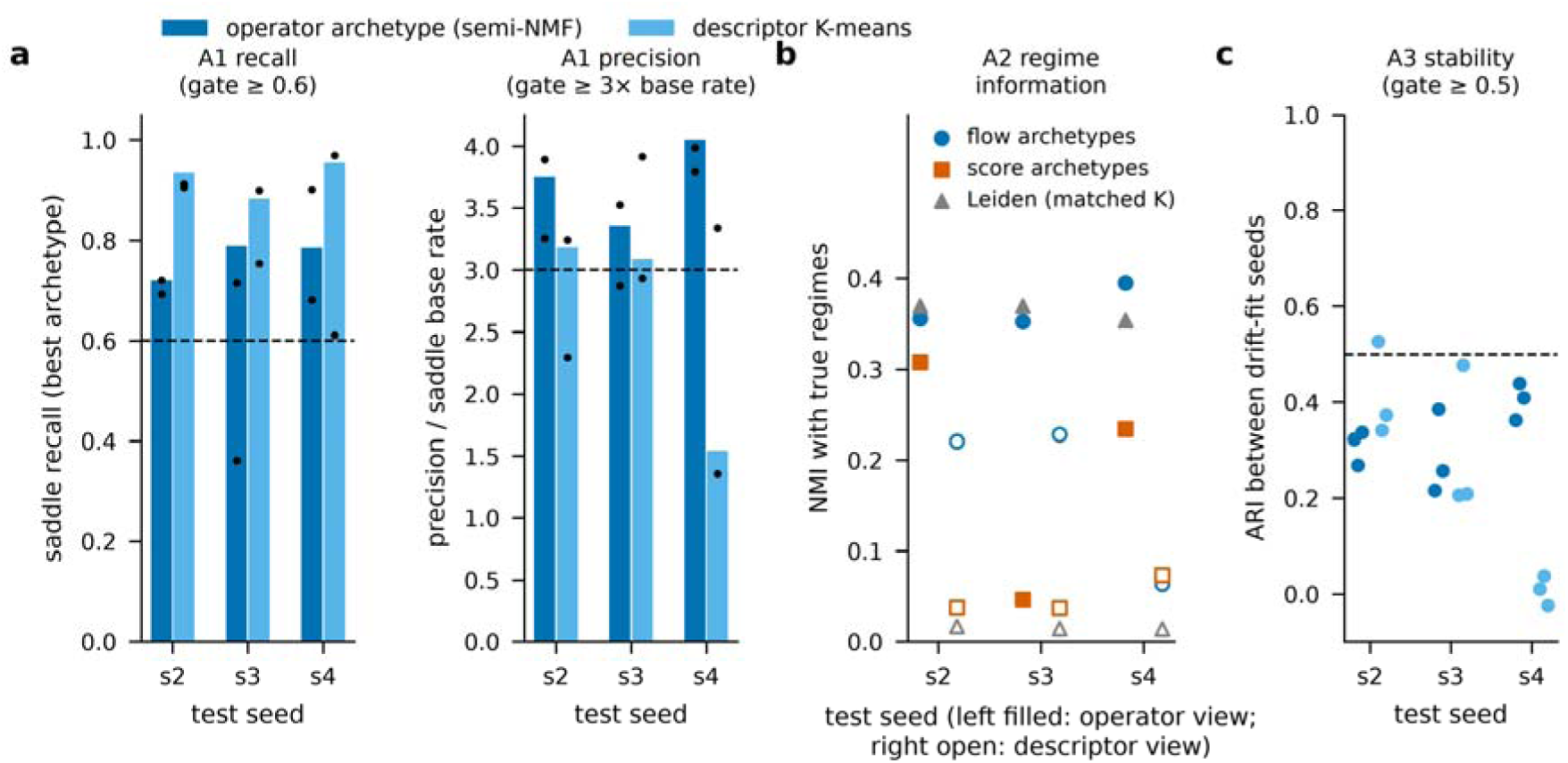
Archetypes of dynamics (S15; pre-registered gates A1–A3). Archetypes were fitted to unit-norm flow operators (semi-NMF, K by held-out reconstruction; operator-archetype view, K = 8) and by K-means on rank-transformed magnitude-free descriptors (descriptor view, K = 2), on the three held-out v3_transition test seeds (s2–s4; PCA-15; S14 reference flow J), and compared with score-operator archetypes and Leiden clustering of expression at matched K. **a**, A1: recall of true saddle cells by the best-matching archetype and its precision relative to the saddle base rate (bars: drift seed 0, the gate rows; dots: drift seeds 1–2, report only). Operator view: recall 0.72–0.79, precision 3.4–4.1× base rate (A1 passes); descriptor view: recall 0.88–0.95, precision 1.5–3.2× (fails on one seed). Dashed lines: gate thresholds (recall ≥ 0.6, precision ≥ 3× base rate). **b**, A2: normalized mutual information with all true regimes. Operator view: flow 0.35–0.39, Leiden 0.35–0.37, score 0.05–0.31; flow beats score on 3 of 3 seeds and Leiden on 1 of 3 (A2 fails). Descriptor view: flow 0.06–0.23, Leiden 0.014–0.017, score 0.04–0.07. **c**, A3: adjusted Rand index of archetype assignments between drift-fit seeds 0, 1, 2 (9 pairs per view): operator view 0.22–0.44, descriptor view −0.02 to 0.53 (gate ≥ 0.5 for every pair; fails). Overall: no view satisfies A1–A3 together.

No view satisfied all three gates. Flow archetypes identify the branching regime specifically, but as a whole partition they carry about as much regime information as expression clusters, and they depend on the fit seed. We report archetypes as a descriptive layer, not as a validated regime classifier.

### 8. Earlier scJDO v1 results

Results of the earlier scJDO v1 analysis — operator handoffs during iPSC reprogramming, branch-specific sensitivity in hematopoiesis, partial concordance with Dynamo and differences from SpliceJAC, and Schrödinger-bridge feasibility in K562 Perturb-seq — are reported in version 56 of this preprint [1]. They used a DSM drift fitted on pseudotime, a score-type operator that the tests above do not validate as dynamics, and are not re-evaluated here.

## Discussion

The useful operator in this study is the flow operator obtained from a measured velocity, and a sampled time course alone is enough to obtain one. Its value is empirical and comparative: on unseen synthetic data it ranked bifurcation cells above committed cells without tuning, on an independent simulator it did so on two of three bifurcations and never reported instability on linear trajectories, and on LARRY its velocity and drift predicted clonal displacement beyond population movement and a held-out day, and it scaled to the full dataset. Lineage tested the velocity, not λ□. It did not recover the values of the true Jacobian, and per-cell operators from independent velocity sources on real data did not agree. We therefore treat the flow operator’s λ□ as an empirical marker of branching in a stated transport space, not as a measurement of a regulatory Jacobian, and all operator-space descriptors as sign- or rank-based.

Representation is a modeling choice with consequences. The signal is carried by the variance-weighted OT metric: it held in PCA of log-normalized counts with 10–20 components, degraded at 50 components and in scVI, and inverted on Factor Analysis and under whitening, although OT velocities in gene space agreed between PCA and FA. Equal-variance representations let noise directions dominate the transport cost. v56 recommended FA for operator inference on the basis of a DSM-drift benchmark on one dataset; that recommendation does not transfer to OT flow operators, and we have corrected it.

Density curvature is not stability out of equilibrium, and in our tests it did not behave as stability even where it was given the best chance. The score operator inverted on non-stationary synthetic presets, did not predict knockdown responses or occupancy changes in K562 and did not predict sister-cell fate coherence in LARRY. These results are consistent with the theory that −log p equals a potential only for a gradient system at steady state [5, 25] and with the known non-identifiability of rotation from snapshots [10]. They are also consistent with v56’s conclusion that the saddle limit of snapshot-fitted drifts is a property of the data modality: in v56, a shuffle test at 100,000 cells showed that the DSM objective was insensitive to the pseudotime input, and on the calibrated benchmark here the operator that ranked saddle cells was the one that used sampling times.

### Relation to existing approaches

The closest prior work falls into five groups, and scJDO2 differs from each in specific, limited ways.

- **ddHodge** reconstructs from RNA velocity a potential, a per-point Hessian of that potential, curl and divergence, and uses divergence as a per-cell stability measure [14]. Its symmetric operator is the Hessian of the gradient part of a velocity field. The score operator here is the Hessian of the log density of a snapshot, needs no velocity and is compared with an independently estimated flow operator per cell.
- **DiffusionOT, TIGON and DeepRUOT** learn a velocity from multi-time-point snapshots by (unbalanced) OT, and DiffusionOT and DeepRUOT also learn a score [3, 4, 29]. TIGON and DiffusionOT differentiate the learned velocity, for gene regulatory networks. Our flow pipeline (entropic OT → barycentric velocity → fitted drift → Jacobian) is an incremental, evaluation-focused variant of this recipe: what we add is per-cell eigen-descriptors, their evaluation against held-out seeds, an independent simulator and lineage barcodes, and the representation analysis.
- **Landscape–flux analyses of scRNA-seq.** Zhu et al. quantify landscape, curl flux, least-action paths and entropy production from an RNA-velocity vector field [15], building on landscape–flux theory [5–7]. Aligning our two operators through a fitted mobility *D* with *f ≈ D□log p* plus flux would apply this framework per cell; for linear systems, the misalignment between drift and diffusion-weighted inverse covariance characterizes non-equilibrium steady states exactly [50]. We have not tested such an alignment and present it as future work, not as a new non-equilibrium measure.
- **Jacobian methods with kinetic velocity.** Dynamo [16], spliceJAC [19] and Barcenas et al. [20] compute Jacobians from splicing or labeling velocity, per cell, per state or along a lineage. When the flow operator takes such velocity as input, it is a Dynamo-style Jacobian, and using λ□ to mark decision cells is the same idea as in spliceJAC, Barcenas et al. and dynamical network biomarker theory [21], applied per cell to an OT-derived drift. Eigenvalue-level identifiability of Jacobian methods was benchmarked by Balubaid et al. [36]. A head-to-head comparison of these methods on a common OT velocity has been run but not scored, and is not reported here.
- **Score-based force decomposition.** Chardès, Maddu and Shelley separate force from intrinsic noise in time-resolved cross-sections by score matching [32, 33], and Boffi and Vanden-Eijnden compute local entropy production from a learned score [34]. These are global field decompositions rather than per-cell operator comparisons, and they are the most direct precedent both for the alignment above and for our finding that diffusion is hard to identify from marginals.

Cell-MNN embeds cells by per-state linear operators learned from time courses [51]; scDiffEq learns neural SDE dynamics and evaluates fate prediction on LARRY [31]; CellRank and CellRank 2 identify macrostates and fate probabilities and accept real-time OT kernels [52, 53]. These answer different questions and are complementary (Table 1).

**Table 1.** Decision guide for single-cell dynamics methods (updated from v56).

| Primary question | Methods | scJDO2 role |
|---|---|---|
| Where will this cell end up? Fate probabilities? | CellRank [52], CellRank 2 [53], Palantir [41] | Complementary; macrostates can annotate operator-space regions |
| How fast are genes changing? | scVelo [18], Dynamo [16] | Input: kinetic velocity for a flow operator, after a velocity-agreement check |
| Which cells at day *t* came from which at day *t − 1*? | Waddington-OT [26], moscot [27] | Input: OT velocity for the label-free flow operator |
| Which cells sit at a branching point, in a time course? | scJDO2 flow operator (λ□), in PCA of log-normalized counts | Ranking only; values not recovered; check representation and scale diagnostics |
| How curved is the cell-state density? | scJDO2 score operator | Descriptive; not a stability measure out of equilibrium |
| When do operator patterns hand off along an observed time axis? | scJDO v1 archetypes [1] | Descriptive, within-pipeline ordering |

### Negative results as part of the method’s specification

We provide the negative results as built-in diagnostics: the representation check (operator read-outs across PCA dimension and against equal-variance representations), the transport-agreement check (mini-batch vs dense OT), the scale check and the null presets (no-fork, linear and stationary). The failures reported here define the conditions under which a flow-operator read-out should not be interpreted.

### Tested and not supported, or not shown

- Recovery of Jacobian values, for either operator (relative error ≈ 1.0).
- Critical-transition timing: two confirmatory tests could not be constructed under the pre-registered eligibility rules, and no model was fitted; the post hoc separation is exploratory.
- Per-cell reproducibility of flow operators across independent velocity sources on real data (null-level agreement on LARRY).
- Diffusion estimation: count noise in the embedding exceeds 48-h diffusion (Supplementary Fig. S1).
- Score-operator prediction of knockdown responses (K562, RPE1) or clonal fate coherence (LARRY).
- Fork detection on every independent-simulator bifurcation (2 of 3) and rotation on every cycle (2 of 3).
- Archetypes as a regime classifier better than expression clustering, or stable across fit seeds.
- Representation-independence of the OT flow operator.
- Novelty relative to work not covered by our literature search (search date 2026-10-26).

### Failure modes

#### Equal-variance representations

OT flow operators should not be computed in FA, whitened or high-dimensional coordinates without the representation check; FA inverted the fork ranking.

#### Coarse sampling

With three sampling times on dyngen, few or no cells were sampled at the branch point, rotation was not resolved and spurious positive eigenvalues appeared in early cells. The number and spacing of time points bound what the flow operator can resolve.

#### Disagreeing velocity sources

On labeling data, the velocity direction depended on whether degradation rates were shared across pulses. A flow operator inherits every ambiguity of its velocity.

#### Static perturbation screens

Without a biologically meaningful trajectory, the score operator did not predict knockdown responses (Results 6); perturbation-response operators are a different identification problem.

#### Cell–cell interactions and composition

Both operators model cell-autonomous dynamics; balanced OT does not model proliferation or death, and fate governed by signaling between cells is outside the model.

### Limitations

scJDO2 estimates local linearized descriptors of a fitted drift or a fitted density in a latent space; neither is a measurement of physical regulatory interactions or a causal gene regulatory network. The synthetic benchmark is calibrated but met only 6 of 13 pre-fixed calibration targets per seed; dyngen is one independent simulator; LARRY is the only lineage reference and scNT-seq hematopoiesis the only labeling reference. Most pre-registered gates failed; the positive findings rest on held-out comparisons of the a-priori configuration and on gates that passed (scaling; flow archetype recall), and they should be read with that in mind. The literature search behind our positioning is dated and may miss recent work. Finally, causal conclusions about any gene or regulator require targeted perturbation.

## Methods

### Embedding for flow operators

Counts were library-normalized to the median library size and log1p-transformed; 2,000 highly variable genes were selected (seurat_v3 flavor on counts) and projected by PCA with k = 15 components (k = 10–20 tested), using raw (unwhitened) coordinates [54]. LARRY real-data comparisons in Supplementary Fig. S4 used PCA-30.

### Time-course velocity

Entropic OT (POT ot.sinkhorn [55], ε = 0.05 × median squared distance, balanced marginals) was computed between consecutive sampling times. Each cell’s velocity is (barycentric projection − position)/Δt; the last time point is extrapolated. For large data, the mini-batch variant uses 4,000-cell source batches in which every cell appears in exactly ten batches, with targets sampled without replacement; barycenters are averaged. Low-rank OT used moscot at rank 50 and 200 [27].

### Drift and flow operator

MeasuredDrift fits an MLP (three hidden layers of 96 units, weight decay 1e-4, single member, seed 0 by default; seeds 1–2 for variability) to the measured velocities; the Jacobian is computed by automatic differentiation at every cell. The tuned configuration of the S14 test was whitened PCA-15, ε factor 0.025, extrapolation, MLP 128×128×128, weight decay 1e-3 and a five-member ensemble.

### Score operator

An energy network (96 hidden units, depth 3) was fitted by score matching with an exact Laplacian [37] (1,500 epochs, learning rate 5e-4, batch size 256); the operator *J_score = −β□²E* (β = 1 in the S11/S22 fits) is the autodiff Hessian. Symmetry was checked per fit (maximum asymmetry ≤ 1e-4). λ□ is the largest eigenvalue.

### Descriptors and scoring

λ□ is the eigenvalue with the largest real part. Fork AUC is the AUC of λ□ for saddle versus committed cells, with truth regions defined from the generator state and used only for scoring. Rotational index is max|Im λ|/max|λ|. Operator-space descriptors (S15) are: sign of Re λ□; number of eigenvalues with Re > 0; Re λ□/max|λ|; max|Im λ|/max|λ|; spectral gap/max|λ|; Henrici non-normality index; transient gain; trace/Σ|λ|, each rank-transformed within dataset (raw-magnitude descriptors were computed for report only). The transition profile is the per-bin median λ□ over 12 day-weighted strict-pseudotime bins, and R is its peak-to-median ratio. Comparison with the true Jacobian used *J_ref*, the Jacobian of the generator’s true expected velocity in the same PCA coordinates (kNN neighborhood 15 or 30), verified by finite differences.

### Calibrated synthetic generator

Twenty real-data statistics (S08) were computed on six datasets; the v3 generator (S09) was calibrated against 13 targets fixed in advance, with a stopping rule that v3 did not meet; the authors accepted v3 with the misses documented. Presets: default, transition (pitchfork with controlled critical point), no-fork, batch, stress and stationary cycle, sampled at days 2, 4 and 6 with lineage barcodes, metabolic labeling and spliced/unspliced counts. Truth regimes (scoring only) are defined from the generator’s core coordinates relative to the critical point (cycling, pre-transition, saddle, committed by branch, other).

### Held-out test of the flow operator (S14)

Part A chose k from {5, 10, 15, 20, 30} on development data by a truth-free rule. The rule was amended after development AUCs had been seen (operator seed stability ≥ 0.5 required before lineage agreement counts), giving k* = 15; the amendment used only the locked truth-free table, and test seeds were untouched. Part B varied one component at a time from the reference on two development seeds and combined the best choices (tuned configuration). Part C evaluated both configurations on held-out datasets (v3_transition s2–s4, v3_default s2–s3, v3_batch s0–s2, v3_stress s0–s2 and v3_nofork s0–s2) at k = 10, 15 and 20, and on LARRY at k = 15. Gates G1–G4 applied to the tuned configuration and the development sanity check; the reference configuration was evaluated in the same blind test.

### Transition eligibility (S14b, S14c)

For each reserve seed, before any flow fit, the truth-only profile (per-bin median of the generator’s critical coordinate) was computed. A dataset was eligible if (a) strict-root provenance was complete, (b) every bin had ≥ 50 cells, (c) the true peak bin was in bins 2–9 and (d) the argmax matched the true bin in ≥ 90 % of 200 bootstrap resamples within bins. Four eligible transitions were required; otherwise the task stopped without fitting. S14b used reserve seeds 5–16, and S14c seeds 17–40 with a 12 GB generation memory guard.

### Independent simulator

dyngen 1.1.1 with backbones bifurcating, cycle and linear (three datasets each), 5,000 cells, 1,000 genes and 64 simulations for bifurcating backbones; synchronized experiments with 8 (S21b) or 3 (S21) time points were re-sampled from saved simulations. Truth was read only after all fits.

### LARRY

GEO GSE140802 [40]; 130,887 cells. Dense analyses used a fixed sample of 4,000 cells per time point (default_rng(0)); barcodes were used for scoring only. Lineage velocity was defined from clonal displacement between sampling days as specified in task S11. Lineage agreement (pre-registered in S14) is the Spearman correlation, pooled over PCA coordinates, between the displacement predicted by integrating the drift from each barcoded day-*t* cell over the sampling gap (1-h Euler steps) and the observed displacement to the centroid of its clone’s day-(*t* + 2) cells; the null permutes cells. The pre-registered test used five permutations; post hoc we used 1,000 and added the controls in Results 5 (population mean-shift predictor, per-interval centring, OT displacement without a drift, magnitudes, λ□ vs displacement magnitude). Bootstrap intervals for held-out energy distances resample day-4 and day-6 cells (200 resamples). Held-out-day prediction fits transport and drift on days 2 and 4 only, integrates the drift from day-4 cells over the 48-h gap (1-h Euler steps) and compares the result with observed day-6 cells by energy distance in a common PCA-50 space, against mean-displacement and no-movement baselines. Day-6 cells are excluded from transport and drift fitting; the gene selection and PCA were fitted on the 12,000-cell sample of all three days, so day 6 contributes to the coordinate system but not to any velocity or drift.

### Scaling (S20)

Dense, mini-batch and low-rank OT were run at 2,000–16,000 cells per day and, for mini-batch OT, on all cells; wall time and peak resident memory were recorded per stage. Agreement with dense OT was measured by median velocity cosine and λ□ Spearman. Cell-type λ□ on all cells is the cell-count-weighted mean of per-day medians.

### Archetypes (S15)

Per-cell operators of the reference flow operator (k = 15) and the score operator were normalized to unit Frobenius norm and decomposed by semi-NMF (ArchetypeDecomposer) with K chosen by held-out reconstruction over {2, 3, 4, 5, 6, 8}; each cell was assigned its dominant archetype. The descriptor view used K-means on rank descriptors with K by mean silhouette over five restarts. Leiden clustering on expression PCA-15 was searched for the matched K. Scores were NMI and ARI with truth regimes, best-matching archetype recall and precision for saddle cells, and ARI across drift seeds 0–2. Gates: A1 (an archetype captures saddle cells with recall ≥ 0.6 and precision ≥ 3× base rate on every test seed), A2 (flow NMI exceeds score and Leiden NMI on every test seed), A3 (ARI ≥ 0.5 across drift seeds); either view could pass.

### Perturbation tests (S18, S18b, S18d)

Replogle K562 and RPE1 CRISPRi Perturb-seq [46] were log-normalized (median library size, log1p, 2,000 control seurat_v3 HVGs, control-only PCA-20). Targets with ≥ 100 cells were tested against controls by energy distance with BH-adjusted permutation p-values. S18 measured the fraction of each significant knockdown’s centroid shift captured by three directions (score-operator soft axes, local or global; control covariance; random subspaces). S18b measured the fraction of each knockdown’s energy distance explained by re-weighting control states (k = 20) and the partial Spearman correlation between score-operator basin stiffness and occupancy sensitivity across basins, with ranks residualized on basin size and covariance trace (10,000 permutations). S18d tested capture of weak-knockdown displacements (lowest knockdown tertile) by bulk-control soft directions in two normalizations (depth-projected log-normalization and analytic Pearson residuals [56]), with RPE1 primary and K562 report only.

### Clonal fate coherence (S22)

The score operator was fitted on LARRY PCA-30 (4,000 cells per day; seeds 0–2, gates on seed 0) and evaluated at every barcoded day-2 and day-4 cell. For clones with ≥ 2 cells on day *t* and ≥ 3 cells on day *t* + 2, the predictor was the median λ□ of day-*t* cells and the truth the Shannon entropy of the day-(*t* + 2) cell-type distribution. Baselines were local density and distance to the day-2 undifferentiated centroid. Spearman correlations were computed pooled, per day and within undifferentiated clones, with 1,000 within-stratum permutations.

### Diffusion (S10)

The effective diffusion trace(D_eff)/d was estimated by clone dispersion and OT noise (Δt = 48 h) and by a score–flow estimator (6 h), and compared with the truth from 32 forward paths per cell; the measurement-noise covariance in the embedding was estimated by redrawing counts from the true pools.

### Pre-registration and review

Every task fixed its data, settings, gates and stopping rules in a written specification before execution (coord/orient/tasks/). Executed tasks were reviewed, and reviews, amendments and user decisions were recorded with dates (coord/orient/reviews/). Amendments made after any result was seen are disclosed where they occur (S14 k rule; v3 acceptance). Each number in this paper is labeled in the figure legends as *pre-registered* (gate fixed in the task before execution), *report only* (fixed in the task but not gated) or *post hoc* (chosen after results). Numbers were recomputed from committed result files by a provenance notebook, which lists the source file and filter for each.

### Statistics

AUCs are rank-based (Mann–Whitney) areas under the ROC curve for λ□ of saddle versus committed cells. Correlations are Spearman unless stated. Permutation nulls: five permutations for the pre-registered LARRY lineage test (1,000 post hoc) and for S11 operator agreement; with five permutations the smallest attainable p is 1/6, so the maximum over the five is reported as a descriptive comparison only; 1,000 within-stratum permutations (S22), 10,000 (S18b) and BH-adjusted permutation tests for perturbation significance. Energy distance between point clouds X and Y in the embedding was E(X, Y) = 2 E□X − Y□ − E□X − X′□ − E□Y − Y′□ [57]. Clustering agreement used NMI and ARI; bootstrap stability used 200 resamples. No correction for multiple comparisons was applied across gates; each gate is a pre-specified pass/fail criterion, and failed gates are reported as results.

### Software

scJDO2 (Python; PyTorch for automatic differentiation). Runtime versions for S14b/S14c: Python 3.13.2, NumPy 2.5.4, pandas 2.3.3, AnnData 0.11.4, Scanpy 1.12.4, Palantir 1.4.5, SciPy 1.18.1, scikit-learn 1.9.1, scvelo 0.3.4, PyTorch 2.14.1; versions for other tasks are recorded in each task report.

## Data Availability

New in v57: LARRY in vitro hematopoiesis (GEO GSE140802) [40]; Replogle et al. 2022 K562 and RPE1 CRISPRi Perturb-seq [46] (figshare article 20029387; files 35773219 for K562 and 35775606 for RPE1); scNT-seq hematopoiesis (GEO GSE193517) [16] and neurons (GEO GSE141851) [39], pancreas (GEO GSE132188) [42] and dentate gyrus (GEO GSE95753) [43] (real-data reference statistics); synthetic generator v3 datasets are regenerated from the manifests and seeds in benchmarks/ (H5AD files are not committed); dyngen [44] settings, seeds and checksums are recorded in benchmarks/dyngen/MANIFEST.csv and the S19/S21/S21b reports; data are regenerated from them.

Datasets used only in the scJDO v1 analyses are listed in version 56 of this preprint.

## Code Availability

scJDO2: https://github.com/manarai/scJDO2 (this version: release v0.6.0 of branch v57-revision, archived on Zenodo, doi:10.5281/zenodo.23699152). The flow operator in the reference configuration is available as one call, scjdo2.flow(adata, time_key=…) (OT velocity, drift, Jacobian and operator-space descriptors; mini-batch transport for large time points). All v57 analyses are in scripts/, with task specifications, reports and reviews in coord/orient/, and number provenance in manuscript/NUMBERS_v57.md. The v56 package and analyses remain available at https://github.com/manarai/scJDO, fixed at git tag scjdo-v56-submission-20260922.

## Author Contributions

Using the CRediT framework:

**David Redd**: Conceptualization, Methodology, Software, Formal analysis, Investigation, Writing — original draft, Writing — review & editing.

**Sam Green**: Methodology, Software, Validation, Data curation, Visualization, Writing — review & editing.

**Tommy W. Terooatea**: Conceptualization, Supervision, Project administration, Funding acquisition, Writing — review & editing.

## Funding

This work was supported by a New Faculty Start-up award from the College of Life Sciences, Brigham Young University (to T.W.T.).

## Competing Interests

The authors declare no competing interests.

## Figure legends

Labels: *pre-registered* = gate fixed in the task before execution; *report only* = fixed in the task but not gated; *post hoc* = chosen after results.

## Supplementary figures

**Supplementary Figure S1.**
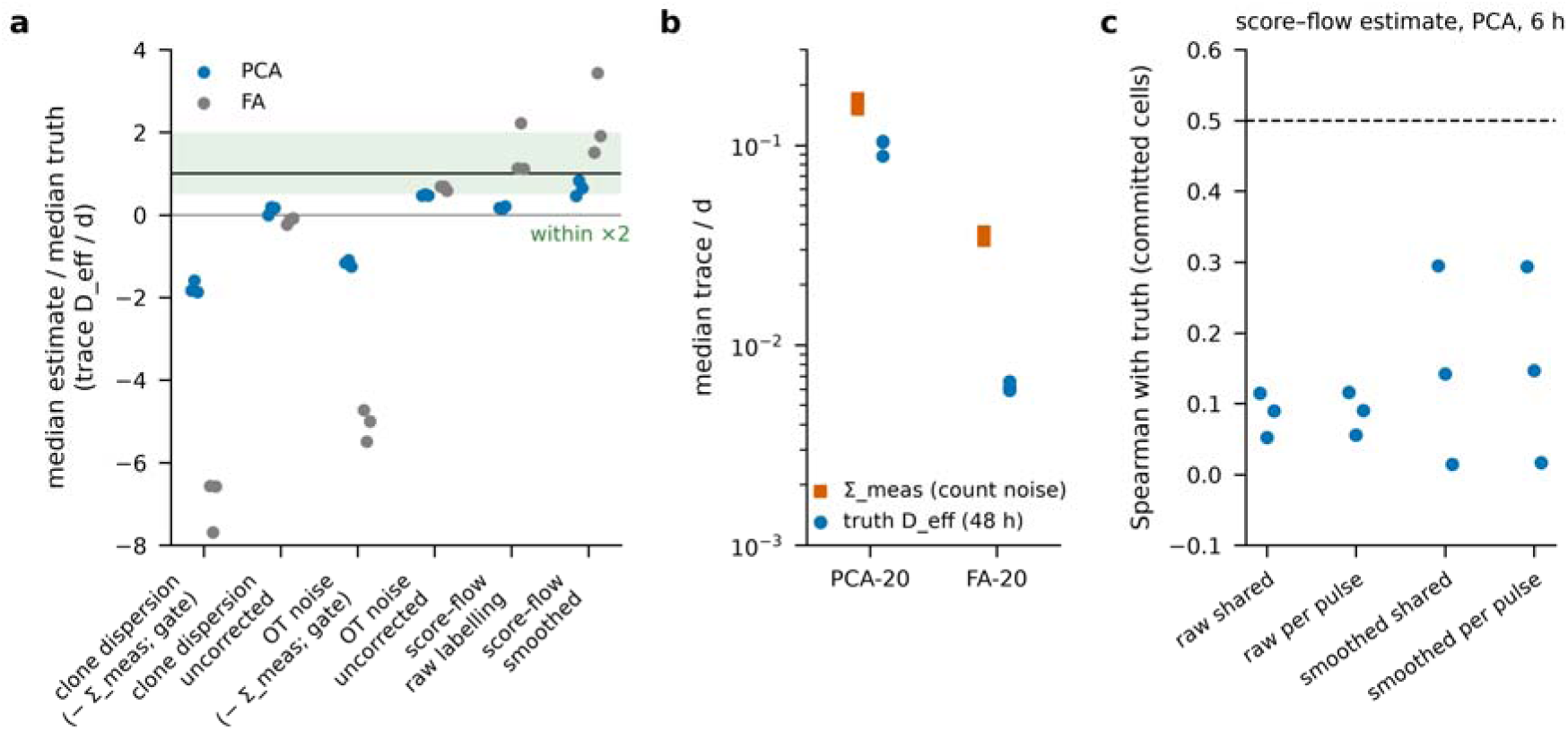
Diffusion is not identifiable at typical sampling intervals (S10, pre-registered; gate failed). v3_default, seeds 0–2. **a**, Median estimate / median truth of trace(D_eff)/d (truth from 32 forward paths per cell, Δt = 48 h for clone and OT estimators, 6 h for the score–flow estimator). Gate versions (Σ_meas subtracted as written): clone dispersion −1.88 to −1.59 (PCA), OT noise −1.25 to −1.09; uncorrected: clone −0.01 to 0.18, OT 0.47–0.50 (the OT held-out selection hit the smallest grid value); score–flow (raw labeling) 0.14–0.20. Green band: within a factor of 2. **b**, Count-noise covariance in the embedding (Σ_meas, counts redrawn from the true pools) exceeds the 48-h effective diffusion: PCA 0.152–0.171 vs 0.088–0.105; FA 0.033–0.037 vs 0.0059–0.0066. **c**, Spearman between the score–flow estimate and the truth over committed cells (gate ≥ 0.5): raw shared γ 0.052–0.115; smoothed shared γ 0.015–0.295.

**Supplementary Figure S2.**
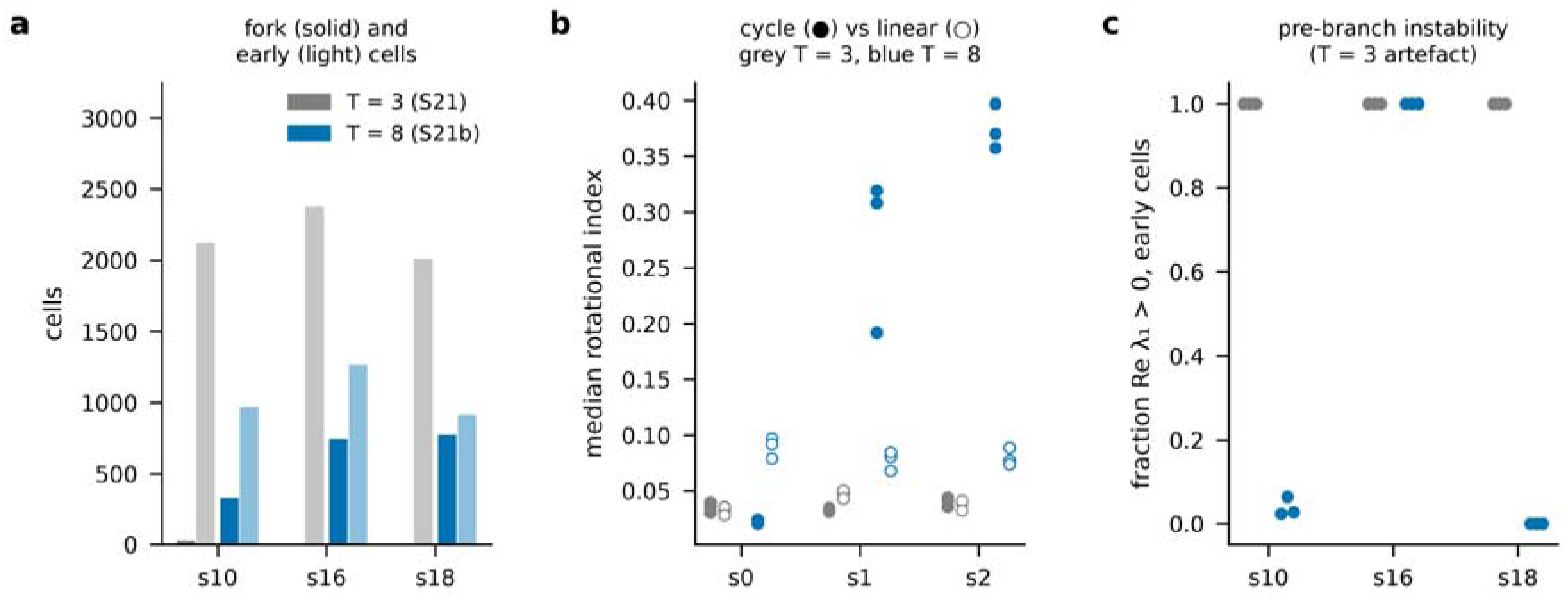
Three sampling times are too coarse on dyngen (S21, pre-registered; both gates failed or not evaluable) vs eight (S21b). Same simulations, synchronized sampling at T = 3 (S21) or T = 8 (S21b); flow J drift seeds 0–2. **a**, Cells in the fork region at T = 3: 22, 0, 0 (s10, s16, s18; F1 not evaluable for s16 and s18) vs 329, 746, 776 at T = 8. **b**, Rotational index, cycle vs linear: cycle above linear on all drift seeds at T = 3 for s0/s1/s2: True, False, True; at T = 8: False, True, True. **c**, Fraction of early (pre-branch) cells with Re λ□ > 0: 1.00 at T = 3 on every dataset and seed; at T = 8, s10 0.02–0.06, s16 1.00, s18 0.00; the S21 reading “pre-branch λ□ > 0” was withdrawn as a sampling artifact.

**Supplementary Figure S3.**
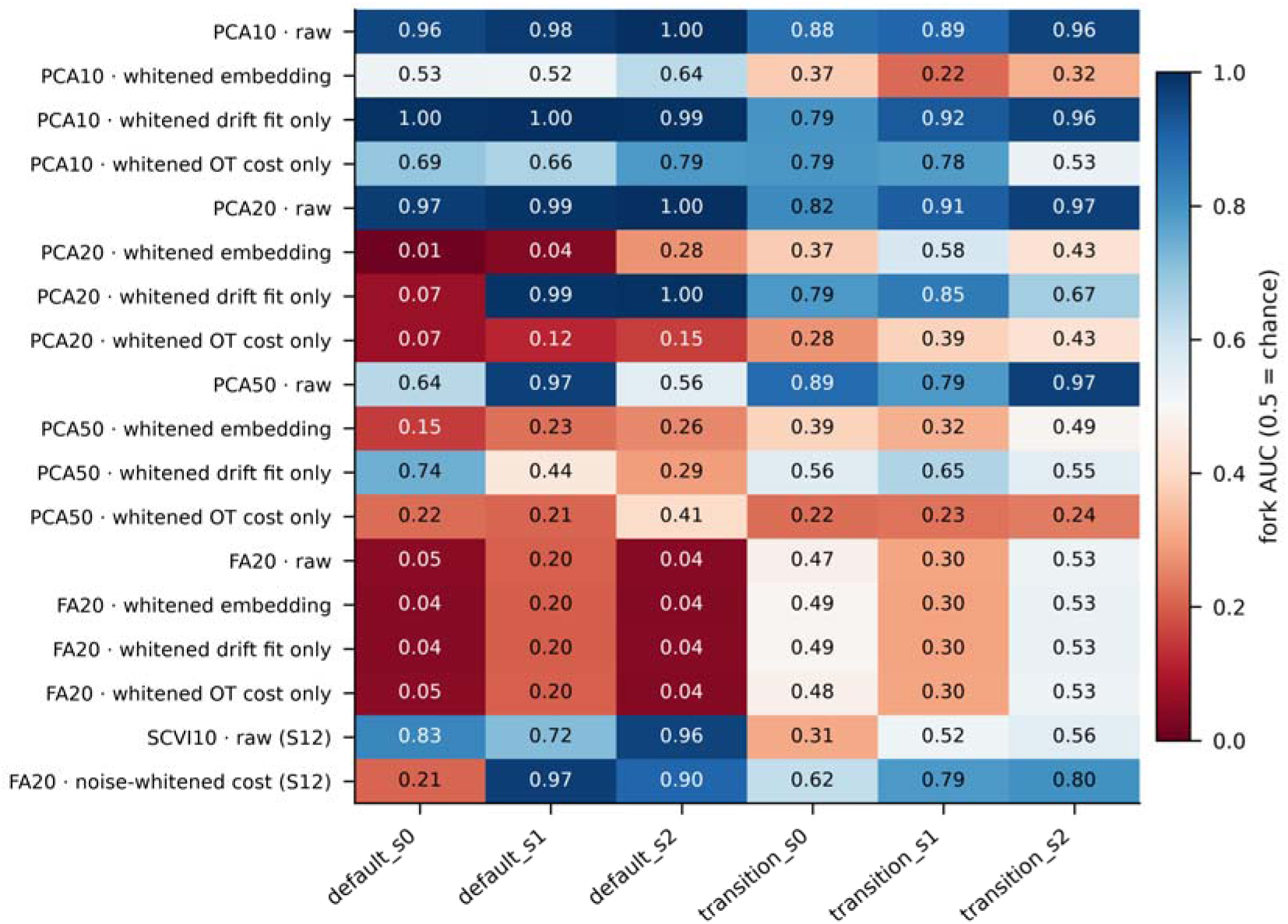
Representation grid for the OT flow operator (S12 Step 0, S13; pre-registered gates failed). Fork AUC (λ□, saddle vs committed) of OT flow J on v3_default and v3_transition seeds 0–2 for PCA-10/20/50 and FA-20, each with four variants: raw coordinates; whitened embedding (OT and drift both in whitened coordinates); whitened drift fit with raw-coordinate OT; whitened OT cost with raw-coordinate drift. Bottom rows: scVI-10 and FA-20 with an OT cost whitened by the FA noise variances (S12 item d). The S13 raw rows reproduce S12 (max |ΔAUC| over PCA-10/20/50/FA-20: 0.0000). Color: blue > 0.5 (saddle cells ranked above committed), red < 0.5 (inverted).

**Supplementary Figure S4.**
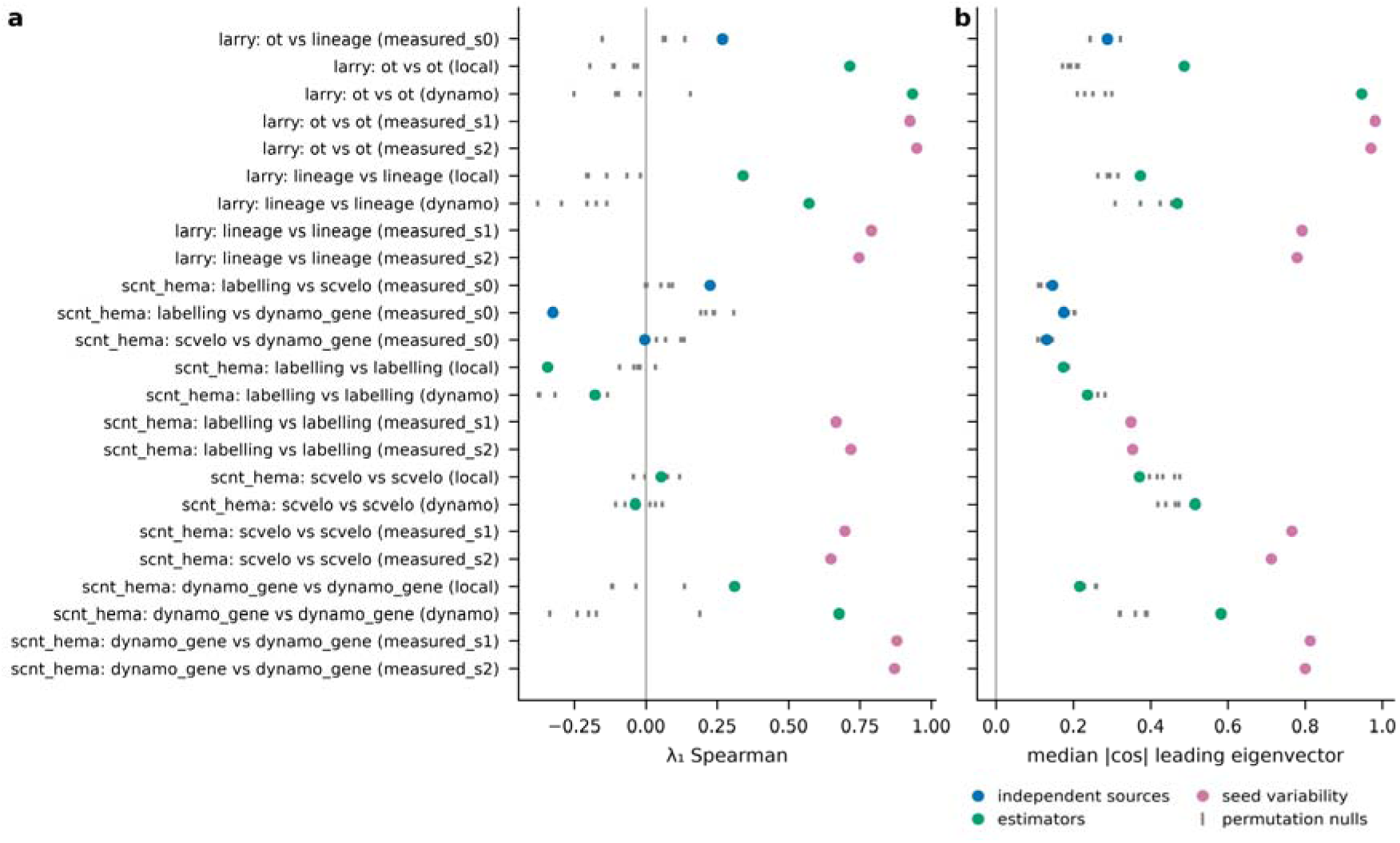
Per-cell flow operators on real data agree across fit seeds and estimators but not across independent velocity sources (S11, pre-registered; 6 of 14 criteria met). LARRY (OT vs lineage velocity, PCA-30, 4,000 cells/day) and scNT-seq hematopoiesis (labeling, scVelo, Dynamo velocities). For each comparison: Spearman of per-cell λ□ (left) and median |cos| of the leading eigenvectors (right); gray ticks: five permutation nulls. LARRY OT vs lineage: λ□ Spearman 0.27 (null max 0.27), |cos| 0.29 (null max 0.32). Seed-variability rows have no null (report only).

## Supplementary tables

- **Supplementary Table S1** — Data sources and accessions (LARRY GSE140802 [40]; scNT-seq hematopoiesis GSE193517 [16] and neurons GSE141851 [39]; Replogle K562/RPE1, figshare 20029387 [46]; scNT-seq hematopoiesis [16]; pancreas GSE132188 [42]; dentate gyrus GSE95753 [43]; dyngen settings [44]).
- **Supplementary Table S2** — S22 clonal fate coherence results (**manuscript/tables/TableS2_S22.csv**): per seed (0–2) and cohort (pooled, Undifferentiated, pooled without tied pairs, day 2 → 4, day 4 → 6), number of clone/time pairs and tied pairs, Spearman ρ of clone λ□ with fate entropy, one-sided permutation p, density and centroid-distance baselines and partial ρ given density; gated rows (seed 0) and gate outcomes (L1–L3 all failed).
- **Supplementary Table S3** — S18d dose-response results (**manuscript/tables/TableS3_S18d.csv**): per screen (RPE1 pre-registered, K562 report only), normalization (N1, N2) and knockdown tertile, numbers of T and M targets, median score-J and covariance capture of T targets, shared-basis null95, criteria D1–D3 and one-sided Wilcoxon and Mann–Whitney p-values; overall outcome underpowered.

## Notes

### Competing Interest Statement

The authors have declared no competing interest.

